# Contrasting effector profiles between bacterial colonisers of kiwifruit reveal redundant roles and interplay converging on PTI-suppression and RIN4

**DOI:** 10.1101/2022.11.12.516272

**Authors:** Jay Jayaraman, Minsoo Yoon, Lauren Hemara, Deborah Bohne, Jibran Tahir, Ronan Chen, Cyril Brendolise, Erik Rikkerink, Matt Templeton

## Abstract

- Testing effector-knockout strains of the *Pseudomonas syringae* pv. *actinidiae* biovar 3 (Psa3) for reduced *in planta* growth in their native kiwifruit host revealed a number of non-redundant effectors that contribute to Psa3 pathogenicity. Conversely, complementation in the weak kiwifruit pathogen *P. syringae* pv. *actinidifoliorum* (Pfm) for increased growth identified redundant Psa3 effectors.
- Psa3 effectors hopAZ1a and HopS2b and the entire exchangeable effector locus (*ΔEEL*; 10 effectors) were significant contributors to bacterial colonisation of the host and were additive in their effects on pathogenicity. Four of the EEL effectors (HopD1a, AvrB2b, HopAW1a, and HopD2a) redundantly contribute to pathogenicity through suppression of pattern-triggered immunity (PTI).
- Important Psa3 effectors include several redundantly required effectors early in the infection process (HopZ5a, HopH1a, AvrPto1b, AvrRpm1a, and HopF1e). These largely target the plant immunity hub, RIN4.
- This comprehensive effector profiling revealed that Psa3 carries robust effector redundancy for a large portion of its effectors, covering a few functions critical to disease.

## Introduction

Bacterial pathogens of plants deploy proteinaceous effectors via their type III secretion system (T3SS) to manipulate their plant hosts and facilitate disease. The *Pseudomonas syringae* species complex delivers as many as 50 secreted effectors to suppress host immunity, as well as to extract nutrients and water from host cells into the apoplastic space (Xin *et al*., 2016; Gentzel *et al*., 2022; Roussin- Léveillée *et al*., 2022). Redundancy within each *P. syringae* strain’s effector repertoire confounds our ability to discern whether particular mechanisms of plant manipulation are universal or host-specific. To date, only the Arabidopsis and tomato pathogen *P. syringae* pv. *tomato* DC3000 (Pto DC3000) has been extensively and comprehensively studied for effector contributions to host infection. Extensive studies in the model plant *Nicotiana benthamiana,* which can also be infected by DC3000 variants, have been particularly important for understanding effector roles in host manipulation (Kvitko *et al*., 2009; Cunnac *et al*., 2011; Wei *et al*., 2015, 2018). However, with largely a single point of pathogen reference, understanding how plant pathogens like *P. syringae* can manipulate many different host plants is challenging.

How pathogens and weak/non-pathogens differ in their colonisation of various host plants is also unclear. The mechanisms of growth within plant hosts for bacterial plant pathogens versus those deployed by the myriad of largely epiphytic commensal bacterial species have only recently been investigated (Chen *et al*., 2020; Velásquez *et al*., 2022). Notably, epiphytic commensal bacteria, much like avirulent pathogenic bacteria that trigger plant immunity, appear to grow only to low but stable numbers *in planta* and display a stationary phase-like growth-death balance (Velásquez *et al*., 2022). A significant proportion of epiphytic commensals, environmentally isolated bacteria, and even symbiotic bacteria possess a functional T3SS but cause little to no disease, thus the role of a T3SS in these species is unclear (Diallo *et al*., 2012; Tampakaki, 2014; Levy *et al*., 2018). The notion of what constitutes a pathogen, including different strategies of colonisation success, may also limit our understanding of the evolution of plant-pathogen relationships (particularly in nature on diverse wild genotypes, as opposed to large human-manipulated plant monocultures). The lessons behind what makes a pathogen versus a commensal strain are critical to understanding how pathogens emerge and what drives their adaptation to cause virulent disease.

The molecular mechanism of plant immunity is currently understood to be comprised of two broad layers: defence at the cell membrane, and intracellular defence. Defence at the plant cell membrane is mediated by transmembrane pattern recognition receptor (PRR) proteins, which recognise evolutionarily conserved pathogen-associated molecular patterns (PAMPs), triggering pattern-triggered immunity (PTI) (DeFalco & Zipfel, 2021). PTI involves a series of plant responses including defence gene expression, hormonal fluxes, apoplastic reactive oxygen species production, and a characteristic callose deposition within the apoplast to block the pathogen incursion (Boller & Felix, 2009; Luna *et al*., 2011). A successful pathogen will overcome PTI through effector deployment. In response to effector presence, plants may deploy their second layer of defence, called effector-triggered immunity (ETI), which is a potentiation and strengthening of PTI responses (Ngou *et al*., 2021; Yuan *et al*., 2021). ETI is triggered intracellularly and is often dependent on effector recognition by polymorphic nucleotide-binding site leucine-rich repeat (NLR) proteins, either directly by binding to effectors, or indirectly through sensing effector presence on guarded proteins: guardees. Often these guardees are protein hubs of PTI or ETI. RPM1-interacting protein 4 (RIN4) is one such immunity hub, is guarded by evolutionarily unlinked resistance proteins in different plants, and is targeted by many different bacterial pathogens (Mackey *et al*., 2002, 2003; Wilton *et al*., 2010; Mazo-Molina *et al*., 2019; Prokchorchik *et al*., 2020; Choi *et al*., 2021).

The kiwifruit bacterial canker pathogen *P. syringae* pv. *actinidiae* (Psa) is a new but growing focus of study for bacterial pathogenesis, in its relationship with its perennial host plant, kiwifruit. Effectors AvrE1d and HopR1b from the particularly virulent Psa biovar 3 (Psa3) have been associated with strong non-redundant contributions to kiwifruit infection (Jayaraman *et al*., 2020). A closely related ubiquitous epiphytic commensal/weak pathogen species, *P. syringae* pv. *actinidifoliorum* (Pfm), has also been described with a functional T3SS and the ability to cause disease on non-kiwifruit plants (Ferrante & Scortichini, 2015; Cunty *et al*., 2015). While several different genetic components have been proposed to be important in woody plant pathogens, Pfm, unlike other epiphytic kiwifruit bacterial colonisers, has all the hallmarks of a successful kiwifruit pathogen: a functional T3SS, a reasonably large repertoire of effectors, and the catechol/β-ketoadipate pathway (Bartoli *et al*., 2015; Nowell *et al*., 2016; Templeton *et al*., 2022). The contrast between Psa3 and Pfm offers an interesting opportunity to study the parameters involved in severe disease outbreaks on plant monocultures, with particular focus on effectors.

## Materials & Methods

### Bioinformatics and sequence analyses

Genome sequences for Psa3 ICMP 18884 (Psa3 V-13; CP011972-3) and Pfm ICMP 18804 (Pfm LV-5; CP081457) were obtained from NCBI GenBank. The Psa3 V-13 and Pfm LV-5 genomes were annotated previously (Templeton *et al*., 2015, 2022). Sequences for type III secreted effectors (T3Es) from Psa3 V-13 and Pfm LV-5 were analysed on Geneious R11 software (https://www.geneious.com; Biomatters) with built-in Geneious DNA and amino acid sequence alignments, tree building, and annotation tools. Effector protein structures were predicted using AlphaFold2 v2.2.0 with a max_template_date of 2022-1-1 (Jumper *et al*., 2021).

### Bacterial strains and growth conditions

The bacterial strains and plasmids used in this study are listed in Supplementary Table S1. Psa3 V-13 and Pfm LV-5 strains were grown in lysogeny broth (LB) at 20°C with shaking at 200 rpm. *Escherichia coli* strains were grown in LB with appropriate antibiotics at 37°C. The concentrations of antibiotics used in selective media were kanamycin 50 μg/mL, gentamicin 25 μg/mL, nitrofurantoin 12.5 μg/mL, cephalexin 40 μg/mL (all from Sigma-Aldrich, Australia). Plasmids were transformed into electrocompetent Psa3 (Mesarich *et al*., 2017) or *E. coli* by electroporation using a Bio-Rad Gene Pulser Xcell and recovered for 1 h in LB before plating on selective media.

### Effector knockout

To make the Pfm LV-5 Δ*hopA1a*, Δ*hopE1a*, or Δ*hopA1a*/Δ*hopE1a* mutants, or Psa3 V-13 Δ*hopH1a*, Δ*hopQ1a*/Δ*hopD1a*, Δ*hopS2b*/Δ*hopAZ1a*, Δ*CEL*/Δ*xEEL*, Δ*CEL*/Δ*hopS2b*/Δ*hopAZ1a*, Δ*CEL*/Δ*xEEL*/Δ*hopS2b*/Δ*hopAZ1a*, Δ*hopH1a*/Δ*hopZ5a*/Δ*avrPto1b*, Δ*hopH1a*/Δ*hopZ5a*/Δ*avrPto1b*/Δ*avrRpm1a* and Δ*hopH1a*/Δ*hopZ5a*/Δ*avrPto1b*/Δ*avrRpm1a*/Δ*tEEL* mutants, methodologies similar to that used for Psa3 V-13 multi-effector knockouts described earlier were used (Hemara *et al*., 2022). Briefly, for each multi-effector knockout, a selected Psa3 V-13 or Pfm LV-5 strain was transformed by electroporation with the relevant pΔ(T3E) construct and transconjugants were selected on LB plates with nitrofurantoin, cephalexin, and kanamycin. Selected colonies were subsequently streaked onto LB plates containing 10% (w/v) sucrose to counter select plasmid integration. Effector mutants were screened using colony PCR with primers Psa_(T3E)- KO_Check-F and Psa_(T3E)-KO_Check-R, and sent for Sanger sequencing with the cloning Psa_(T3E)- KO_UP-F and Psa_(T3E)-KO_DN-R primers described earlier (Hemara *et al*., 2022). Mutants were also confirmed by plating on kanamycin-containing medium to confirm loss of the integrated *nptII* gene (and associated *sacB* gene).

### Effector plasmid complementation

For native-promoter constructs of Pfm LV-5 effectors, the full region including the HrpL box promoter was PCR-amplified using primers (Supplementary Table S2) and Q5 High-Fidelity DNA Polymerase (NEB, USA). The resulting PCR fragment was gel-purified and was blunt-end-ligated into the *Eco53k*I (NEB) site of broad host-range vector pBBR1MCS-5 (Kovach *et al*., 1995). Constructs were transformed into *E. coli* DH5α, plated on X-gal/IPTG-containing (for blue/white selection) LB agar plates with gentamicin, and positive transformants confirmed by Sanger sequencing (Macrogen, South Korea).

Synthetic *avrRps4* promoter constructs of HA-tagged effectors from Psa3 V-13 or Pfm LV-5 have been described previously (Jayaraman *et al*., 2017). All constructs were transformed into relevant Psa3 or Pfm strains by electroporation, and transformants screened for presence of effector by gene-specific colony PCR.

### *In planta* growth and symptomology assays

Psa3 and Pfm infection assays were carried out as described previously (McAtee *et al*., 2018). *A. chinensis* var. *chinensis* ‘Hort16A’ plantlets, grown from axillary buds on Murashige and Skoog rooting medium without antibiotics in sterile 400-mL plastic tubs (“pottles”), were purchased from Multiflora (Auckland, New Zealand). Plantlets were grown at 20°C under Gro-Lux fluorescent lights under long-day conditions (16 h:8 h, light:dark) and used when the plantlets were approximately 12 weeks old. Overnight LB medium cultures of Psa3 or Pfm were pelleted at 5,000*g*, resuspended in 10 mM MgSO_4_, reconstituted at OD_600_ = 0.05 (c. 10^6^ cfu/mL, determined by plating) in 500 mL of 10 mM MgSO_4_. Surfactant Silwet L-77 (Lehle Seeds, TX, USA) was added to the inoculum at 0.0025% (vol/vol) to facilitate leaf wetting. Pottles of ‘Hort16A’ plantlets were flooded with the inoculum, submerging the plantlets for 3 min, drained, sealed, and then incubated under plant growth conditions, as above.

*In planta* growth of Psa3 or Pfm strains was assayed as described previously (McAtee *et al*., 2018). Briefly, leaf samples of four leaf discs per pseudobiological replicate, taken randomly with a 1-cm diameter cork-borer from three plants, were harvested at 2 h (day 0), day 6, and day 12 post-inoculation. All four replicates per treatment, per time point were taken from the same pottle. To determine Psa3/Pfm growth inside the plant, the leaf discs were surface-sterilised, placed in Eppendorf tubes containing three sterile stainless-steel ball bearings, 350 μL 10 mM MgSO_4_, and macerated in a Storm 24 Bullet Blender (Next Advance, NY, USA) for two bursts of 1 min each at maximum speed. A 10-fold dilution series of the leaf homogenates was made in sterile 10 mM MgSO_4_ until a dilution of 10^−8^ and plated as 10 μL droplets on LB medium supplemented with nitrofurantoin and cephalexin. After 2 days of incubation at 20°C, the cfu per cm^2^ of leaf area was ascertained from dilutions. To observe pathogenic symptoms on the plants, infected pottles were kept up to 50 days post-inoculation and photographs taken of pottles and a representative infected leaf. Infection severity was qualitatively assessed based on typical symptoms: necrotic leaf spots, chlorotic haloes, leaf death, and plant death. Each of these growth assay experiments was conducted at least three times.

### PTI-suppression assays

The *N. benthamiana* PTI-suppression assay (suppression of effector delivery) was adapted from that described previously (Crabill *et al*., 2010; Le Roux *et al*., 2015). pBBR1MCS-5 constructs of each Psa3 V-13 or Pfm LV-5 effector (Jayaraman *et al*., 2017) were transformed by electroporation into Pfo Pf0- 1 (T3S) strains (Thomas *et al*., 2009) and plated on selective media with chloramphenicol, gentamicin, and tetracycline. Positive transformants were confirmed by gene-specific colony PCR. Pf0-1(T3S) carrying empty vector or Psa3/Pfm constructs were streaked from glycerol stocks onto LB agar plates with antibiotic selection and grown for 2 days at 28°C. Bacteria were then harvested from plates, resuspended in 10 mM MgSO_4_, and diluted to the required OD_600_ = 0.6 (c. 10^9^ cfu/mL). Infiltrations were carried out on fully expanded leaves of 4- to 5-week-old *Nicotiana benthamiana* using a blunt-end syringe on two or three leaves (replicates). Next, 12 hours post-infiltration, Pto DC3000 (OD_600_ = 0.03; c. 10^7^ cfu/mL) was infiltrated in an overlapping area of the leaves. Pto DC3000-triggered tissue collapse was scored at 3 dpi. PTI suppression experiments were conducted in triplicate, over three independent experimental runs, with tissue collapse in at least 50% of replicates scored as suppressors of PTI.

The *A. chinensis* PTI-suppression assay (suppression of callose deposition) was adapted from that described previously (Jin & Mackey, 2017). Briefly, for observation of callose deposits, Pfo Pf0-1 (T3S) carrying either empty vector or the plasmid-borne Psa3 effector (as before) was vacuum-infiltrated into *A. chinensis* leaves from plantlets grown in tissue culture (Multiflora, NZ) at 10^8^ cfu/mL (OD_600_ of 1 in sterile 10mM MgSO_4_). The infected leaves were decolorised in lactophenol solution (water 8.3%, glycerol 8.3%, lactic acid 7%, water saturated phenol 8.3%; in ethanol v/v) and then stained with 0.01% aniline blue in 150 mM K_2_HPO_4_, pH 9.5 (all chemicals from Sigma Aldrich). Callose deposits were visualised with a Nikon Ni-E upright compound UV-fluorescence microscope equipped with a digital camera under a 40x magnification, and acquired images analyzed using ImageJ software by determining the average area of a single callose deposit and then calculated callose counts based on total callose deposit area in each image.

### *In vitro* effector secretion assay

For detection of effector secretion *in vitro*, the protocols used were based on those described previously (Huynh *et al*., 1989). Briefly, Psa3 V-13 or Pfm LV-5 strains carrying the relevant HA-tagged effector plasmid constructs (pBBR1MCS-5) were grown in LB medium with antibiotic selection overnight, pelleted at 5,000*g*, washed with *hrp*-inducing minimal medium supplemented with 10 mM fructose and then resuspended in *hrp*-inducing minimal medium and incubated for 6 h with shaking for *hrp* induction. Following *hrp* induction, cells were pelleted and proteins extracted by the Laemmli method (Laemmli, 1970), resolved by SDS-PAGE and immunoblotted for the presence of the HA-tagged effector using α-HA antibody (H9658; Sigma-Aldrich) and α-HA-HRP (3F10; Roche, Basel, Switzerland).

### Reporter eclipse assay

Freshly expanded leaves of *A. chinensis* var. *chinensis* ‘Hort16A’ were co-bombarded with DNA-coated gold particles carrying pRT99-GUS and pICH86988 with the effector of interest, as described in Jayaraman et al. (2021). Effectors were YFP-tagged and cloned under a CaMV 35S promoter (Choi *et al*., 2017).

### Transient expression in *Nicotiana benthamiana* and co-immunoprecipitation

*Agrobacterium tumefaciens* AGL1 (YFP-tagged effectors; (Choi *et al*., 2017)) or GV3101 pMP90 (FLAG-tagged AcRIN4s; (Yoon & Rikkerink, 2020)) was freshly grown in LB with appropriate antibiotics at 28°C with shaking at 200 rpm. Cells were pelleted by centrifugation at 4000 *g* for 10 min and resuspended in infiltration buffer (10 mM MgCl_2_, 5 mM EGTA, 100 μM acetosyringone). Cell suspensions were diluted to a final OD_600_ of 0.1 and infiltrated into at least two fully expanded leaves of 4- to 5-week-old *N. benthamiana* plants using a needleless syringe. All *Agrobacterium*-mediated transformation experiments were performed using pre-mixed *Agrobacterium* cultures for the stipulated effector-RIN4 combinations in a single injection for co-immunoprecipitation experiments (see below). YFP was used as a negative control for effectors.

Tissues (0.5 g per sample) were collected 2 days post-infiltration and ground to a homogeneous powder in liquid nitrogen and resuspended in 1 mL of protein extraction buffer (1× PBS, 1% *n*-dodecyl- β-d-maltoside or DDM (Invitrogen, Carlsbad, CA, USA), and 0.1 tablet cOmplete™ protease inhibitor cocktail (Sigma-Aldrich) in NativePAGE™ buffer (Invitrogen)). Extracted protein samples were centrifuged at 20 000*g* for 2 min at 4°C and the supernatant was collected for immunoprecipitation using the µMACS GFP Isolation Kit (Miltenyi Biotec, MediRay, New Zealand). Total and immunoprecipitated proteins were resolved on a 4–12% SDS-PAGE gel. Western blots using PVDF membranes were prepared and probed using HRP-conjugated antibodies in 0.2% I-Block (Invitrogen).

Detection was achieved using ECL (Amersham, GE Healthcare, Chicago, IL, USA). The antibodies used were α-FLAG (F1804; Sigma-Aldrich), α-FLAG-HRP (A8592; Sigma-Aldrich), and α-GFP (MA515256; Life Technologies).

## Results

### Three new effector loci contribute quantitatively and additively to Psa3 pathogenicity

When comparing their capacity for virulence, Pfm LV-5 is clearly incapable of causing the prolific disease symptoms in ‘Hort16A’ that Psa3 V-13 can, despite both species being commonly recovered from kiwifruit plants in the orchard (Figure 1A) (Chapman *et al*., 2012; Vanneste *et al*., 2013; McCann *et al*., 2013; Cunty *et al*., 2015; Abelleira *et al*., 2015). Comparing the pathogenicity of Pfm LV-5 to Psa3 V-13 and Psa3 V-13 carrying the avirulence effector *hopA1j* from *P. syringae* pv. *syringae* 61 indicated that Pfm LV-5 more closely resembled the avirulent strain than pathogenic wild-type Psa3 V-13 (Figure 1B).

**Fig. 1.**
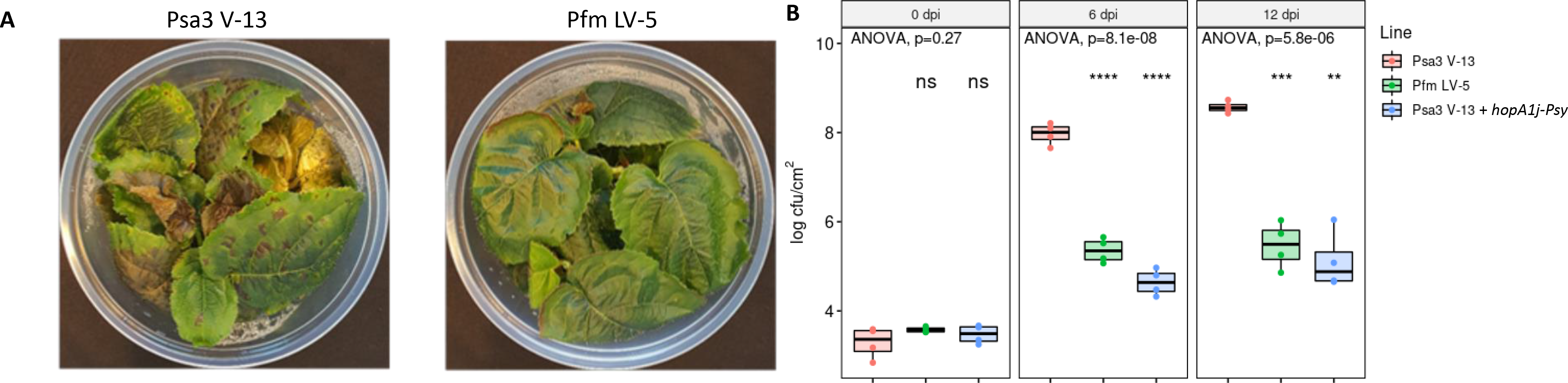
*Pseudomonas syringae* pv. *actinidifoliorum* (Pfm) LV-5 lacks pathogenicity in kiwifruit plants in comparison to *P. syringae* pv. *actinidiae* biovar 3 (Psa3) V-13. (A) *Actinidia chinensis* var. *chinensis* ‘Hort16A’ plantlets were flood inoculated with wild-type Psa3 V-13 or Pfm LV-5 at approximately 10_6_ cfu/mL. Photographs of symptom development on representative pottles of ‘Hort16A’ plantlets at 50 days post-infection. (B) ‘Hort16A’ plantlets were flood inoculated with wild-type Psa3 V-13, Pfm LV-5, or Psa3 V-13 carrying plasmid-borne avirulence effector *hopA1j* (from *P. syringae* pv. *syringae* 61) at approximately 10 cfu/mL. Bacterial growth was quantified at 6 and 12 days post-inoculation by serial dilution and plate count quantification. Box and whisker plots, with black bars representing the median values and whiskers representing the 1.5 inter-quartile range, for *in planta* bacterial counts plotted as Log_10_ cfu/cm from four pseudobiological replicates. Asterisks indicate statistically significant differences from Student’s t-test between the indicated strain and wild-type Psa3 V-13, where p≤.01 (**), p≤.001 (***), or p≤.0001 (****); not significant (ns). These experiments were conducted three times on independent batches of ‘Hort16A’ plants, with similar results.

Previously, *avrE1d* and *hopR1b* were found to be required for virulence (disease symptoms) and pathogenicity (host colonisation) in Psa3 V-13 infection of ‘Hort16A’ (Jayaraman *et al*., 2020). Pfm LV- 5 carries orthologs of both AvrE1 and HopR1 and these versions share both amino acid identity and predicted protein structure to orthologs in Psa3 V-13 (Figure 2; Supplementary Table S3). Both AvrE1d and HopR1b appear to function non-redundantly as putative pore-forming effectors, a role apparently shared by HopAS1b in multiple pseudomonads (Figure 2). Surprisingly, however, loss of *hopAS1b* has not yet been identified to significantly affect virulence of Psa3 V-13.

**Fig. 2.**
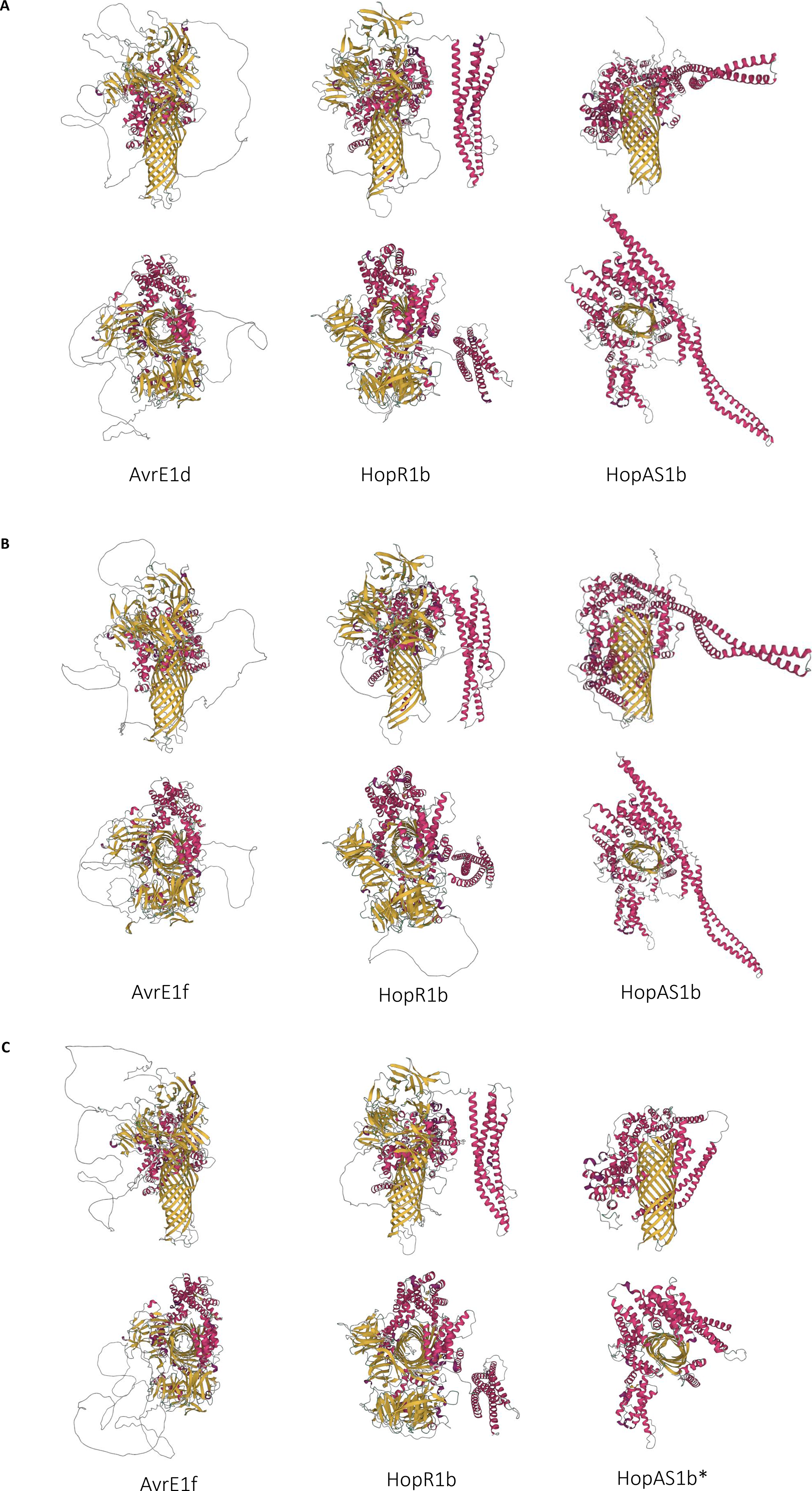
*Pseudomonas syringae* pv. *actinidiae* biovar 3 (Psa3) V-13 and *P. syringae* pv. *actinidifoliorum* (Pfm) LV-5 share multiple pore-forming effector orthologs. Predicted protein structures for AvrE1, HopR1b, and HopAS1b from (A) Psa3 V-13, (B) Pfm LV-5 and (C) Pto DC3000. HopAS1b* is the non-translated C-terminal sequence portion of HopAS1b from Pto DC3000, translated from after the frameshift mutation. This sequence lacks the N-terminal portion before the frameshift mutation and is probably untranslated, since translation is initiated from the HrpL promoter site. Structures were predicted using AlphaFold2. Alpha helices are coloured pink and beta sheets are coloured yellow.

To investigate effector requirements and contribution to virulence and pathogenicity in susceptible *A. chinensis* var. *chinensis* plants, a previously developed library of effector knockout strains was tested on ‘Hort16A’ plantlets and assessed for reduced *in planta* colonisation (Hemara *et al*., 2022). Reductions in Psa3 V-13 growth, assessed at 6 and 12 dpi, were observed in *ΔhopS2a*, *ΔhopAZ1a*, and *ΔxEEL* mutant strains (Figure 3A; Figure S1). Despite having a similar topology to AvrE1d and HopR1b, loss of *hopAS1b* did not affect virulence or pathogenicity of Psa3 V-13. Interestingly, unlike the symptom reduction seen previously for loss of effectors *avrE1d* (*ΔCEL*) and *hopR1b* (Jayaraman *et al*., 2020), loss of these three effector loci was not associated with a reduction in disease symptom progression on the highly susceptible ‘Hort16A’ plants (Figure 3B). Plasmid complementation of effectors *hopS2a* (with its chaperone *shcS2*) and *hopAZ1a* in their respective knockout strains restored host colonisation (Figure S2).

**Fig. 3.**
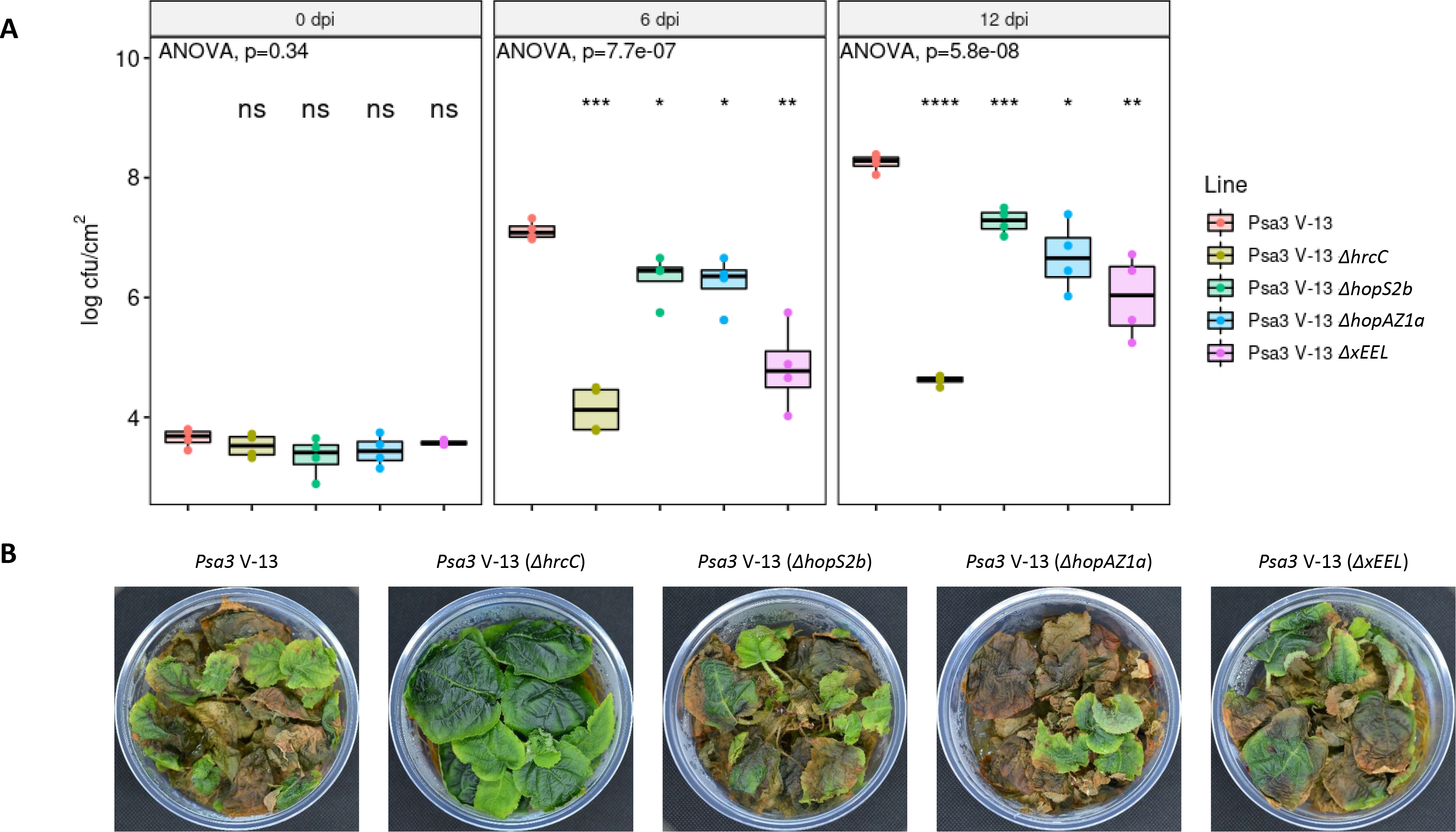
Three *Pseudomonas syringae* pv. *actinidiae* biovar 3 (Psa3) V-13 effector loci are independently required for full pathogenicity but not required for virulence. *Actinidia chinensis* var. *chinensis* ‘Hort16A’ plantlets were flood inoculated with wild-type Psa3 V-13, Δ*hrcC* mutant, Δ*hopS2b* mutant, Δ*hopAZ1a* mutant or Δ*xEEL* (extended EEL) mutant at approximately 10^6^ cfu/mL. (A) Bacterial growth was quantified at 6 and 12 days post-inoculation by serial dilution and plate count quantification. Box and whisker plots, with black bars representing the median values and whiskers representing the 1.5 inter-quartile range, for *in planta* bacterial counts plotted as Log_10_ cfu/cm^2^ from four pseudobiological replicates. Asterisks indicate statistically significant differences from Welch’s t- test between the indicated strain and wild-type Psa3 V-13, where p≤.05 (*), p≤.01 (**), p≤.001 (***), or p≤.0001 (****); not significant (ns). These experiments were conducted three times on independent batches of ‘Hort16A’ plants, with similar results. (B) Symptom development on representative pottles of ‘Hort16A’ plantlets infected with strains in (A) at 50 days post-infection.

The ten-effector *ΔxEEL* knockout mutant resulted in a loss of pathogenicity and contrasted with the eight-effector *ΔfEEL* knockout mutant that remained fully pathogenic (Figure S1). To investigate whether the additional two effectors lost (*ΔhopQ1a* and *ΔhopD1a*) were redundantly responsible for the contribution to pathogenicity in the *ΔxEEL* mutant, a double knockout of these two effectors was generated and tested alongside *ΔfEEL* and *ΔxEEL* mutants (Figure 4A). Notably, neither the *ΔfEEL* nor the *ΔhopQ1a*/ *ΔhopD1a* mutant strains showed reduced pathogenicity, suggesting that loss of effector redundancy across the total set of ten effectors in the x*EEL* locus was probably responsible for the change in the *ΔxEEL* mutant (Figure 4B).

**Fig. 4.**
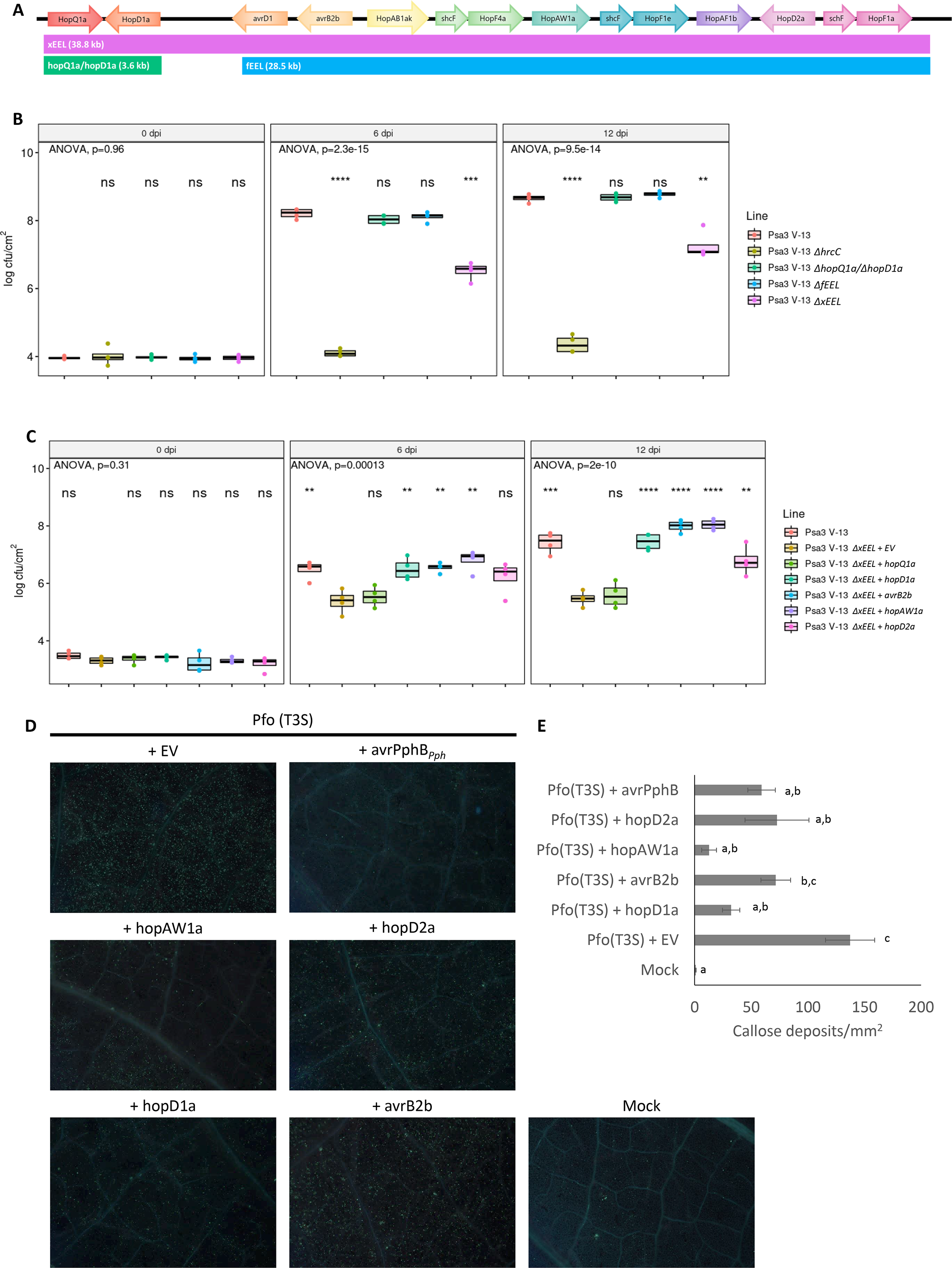
The *Pseudomonas syringae* pv. *actinidiae* biovar 3 (Psa3) V-13 extended exchangeable effector locus (xEEL) carries four redundantly required PTI-suppressing effectors. (A) The extended EEL (xEEL) of Psa3 V-13 is made up of ten effectors (*hopQ1a* to *hopF1a*) with a smaller subset of eight effectors (*avrD1* to *hopF1a*) designated as the full EEL (fEEL). (B) *Actinidia chinensis* var. *chinensis* ‘Hort16A’ plantlets were flood inoculated with wild-type Psa3 V-13, Δ*hrcC* mutant, Δ*hopQ1a*/Δ*hopD1a* double mutant, Δ*fEEL* mutant, or Δ*xEEL* mutant at approximately 10_6_ cfu/mL. Bacterial growth was quantified at 6 and 12 days post-inoculation by serial dilution and plate count quantification. (C) *A. chinensis* var. *chinensis* ‘Hort16A’ plantlets were flood inoculated with wild-type Psa3 V-13 carrying an empty vector (+EV), Δ*xEEL* mutant +EV, or Δ*xEEL* mutant complemented with plasmid vector carrying *hopQ1a*, *hopD1a*, *avrB2b*, *hopAW1a*, or *hopD2a* at approximately 10_6_ cfu/mL. Bacterial growth was quantified at 6 and 12 days post-inoculation by serial dilution and plate count quantification. In (B) and (C), data are presented as box and whisker plots, with black bars representing the median values and whiskers representing the 1.5 inter-quartile range, for *in planta* bacterial counts plotted as Log_10_ cfu/cm_2_ from four pseudobiological replicates. Asterisks indicate statistically significant differences from Welch’s t- test between the indicated strain and Psa3 V-13 Δ*xEEL* mutant carrying empty vector, where p≤.01 (**), p≤.001 (***), or p≤.0001 (****); not significant (ns). These experiments were conducted three times on independent batches of ‘Hort16A’ plants, with similar results. (D) Callose deposition induced by *P. fluorescens* Pf0-1 (T3S) strain carrying empty vector (+ EV), or positive control HopAR1 effector (+ AvrPphB), or one of four Psa3 V-13 effectors from (B) in *A. chinensis* var*. deliciosa* leaves. The representative images were captured at 48 h after infiltration with mock (sterile 10mM MgSO_4_) or bacterial strains. (E) The number of callose deposits per mm^2^ of leaf tissue from (D) was analyzed with the ImageJ software. Mean and standard error (SEM<) were calculated with results from five biological replicates. Different letters indicate significant difference from a one-way ANOVA and Tukey’s HSD *post hoc* test at p≤.05.

Screening all Psa3 V-13 effectors for PTI-suppression activity previously identified HopD1a as a potent contributor to PTI suppression (Crabill *et al*., 2010; Choi *et al*., 2017). Interestingly, using *P. fluorescens* Pf0-1 (T3S) delivery for re-screening effectors from Psa3 V-13, with a lower stringency (suppression was considered positive if at least 2 out of 4 infiltrated leaf patches showed a hypersensitive response), identified four Psa3 effectors that were robustly able to suppress *P. fluorescens*-triggered PTI in *N. benthamiana* plants: HopD1a, AvrB2b, HopD2a, and HopAW1a (Figure S3). All four effectors lie within the *xEEL* locus and were able to suppress PTI to allow for the subsequent ETI triggered by Pto DC3000 to a capacity comparable to that of the positive control, AvrPtoB from Pto DC3000 (Figure S3). Testing of individual effector contributions to the *ΔxEEL* mutant by plasmid complementation confirmed that these four effectors were individually able to restore the *ΔxEEL* mutant’s loss of pathogenicity (Figure 4C). To assess whether these effectors were also able to suppress PTI in their natural plant host, *A. chinensis* leaves were used to assess callose deposition (a PTI response) against *P. fluorescens* Pf0-1 (T3S) carrying empty vector or each of the four *xEEL* effectors (Figure 4D). Notably, three out of the four effectors were able to significantly suppress callose deposition as expected (Figure 4D–4E).

To determine whether the three effectors/loci (*hopS2*, *hopAZ1* and *xEEL*) additively contribute to pathogenicity and virulence in ‘Hort16A’, cumulative knockouts were generated and tested for *in planta* colonisation and symptom development. These effector contributions were tested in the *ΔCEL* background where *avrE1d* contributes a large proportion of the pathogenicity and virulence observed in Psa3 (Jayaraman *et al*., 2020). Knockout of the *xEEL* in the *ΔCEL* background, or knockout of *hopS2b* and *hopAZ1a* in addition to *ΔCEL,* reduced pathogenicity of the double and triple mutant strains, respectively, to a similar extent as that in the avirulent *ΔhrcC* mutant (Figure 5A). Unsurprisingly, knocking out these effectors in addition to the loss of the *CEL* did not show disease progression differences from those seen in the symptomless *ΔCEL*-infected ‘Hort16A’ plants (Figure 5B). Pathogenicity and virulence were also tested for the quadruple-locus knockout of *ΔCEL*/*ΔxEEL*/*ΔhopS2b*/*ΔhopAZ1a* and found to be no different from the *ΔhrcC* mutant either (Figure S4). Taken together, these assays have identified three new effector loci that non-redundantly and additively contribute to virulence and pathogenicity of Psa3 in ‘Hort16A’ plants.

**Fig. 5.**
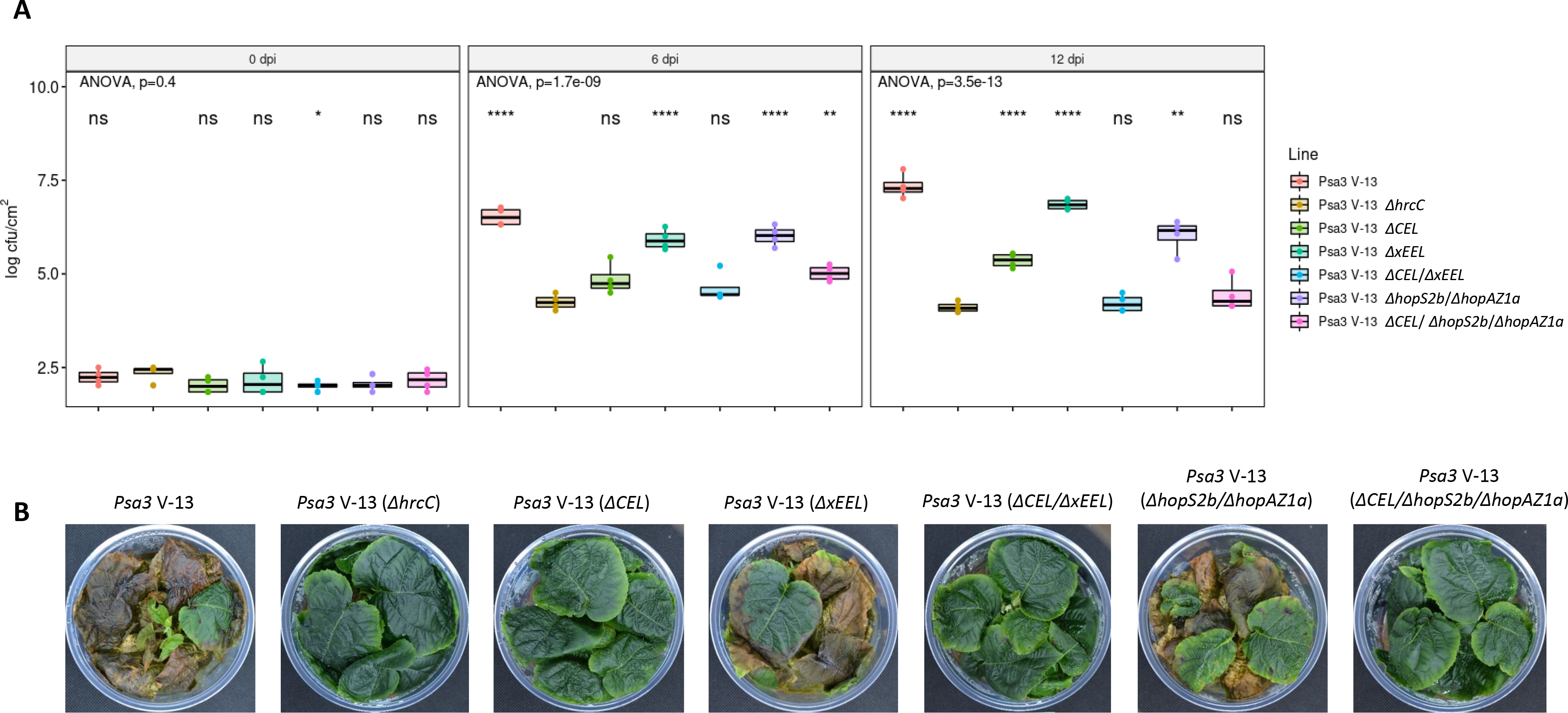
*Pseudomonas syringae* pv. *actinidiae* biovar 3 (Psa3) V-13 pathogenicity-associated effector loci are required alongside the conserved effector locus (CEL) for pathogenicity. *Actinidia chinensis* var. *chinensis* ‘Hort16A’ plantlets were flood inoculated with wild-type Psa3 V-13, Δ*hrcC* mutant, Δ*CEL* mutant, Δ*xEEL* mutant, Δ*CEL*/Δ*xEEL* double mutant, Δ*hopS2b*/Δ*hopAZ1a* double mutant, or Δ*CEL*/Δ*hopS2b*/Δ*hopAZ1a* triple mutant at approximately 10 cfu/mL. (A) Bacterial growth was quantified at 6 and 12 days post-inoculation by serial dilution and plate count quantification. Box and whisker plots, with black bars representing the median values and whiskers representing the 1.5 inter-quartile range, for *in planta* bacterial counts plotted as Log_10_ cfu/cm from four pseudobiological replicates. Asterisks indicate statistically significant differences from Welch’s t-test between the indicated strain and the Psa3 V-13 Δ*hrcC* mutant, where p≤.05 (*), p≤.01 (**), or p≤.0001 (****); not significant (ns). These experiments were conducted three times on independent batches of ‘Hort16A’ plants, with similar results. (B) Symptom development on representative pottles of ‘Hort16A’ plantlets infected with strains in (A) at 50 days post-infection.

### Avirulence effectors from Pfm cannot explain its lack of virulence in ‘Hort16A’

Surprisingly, four effectors (*avrE1d*, *hopR1b*, *hopS2b*, and *hopAZ1a*) identified from Psa3 V-13 that individually contribute significantly to pathogenicity and virulence were also present in Pfm LV-5, along with required promoters and chaperones (Supplementary Table S3). The exception to required effectors in Psa3 also being present in Pfm were effectors from the *xEEL* in Psa3 V-13 (*hopD1a*, *avrB2b*, *hopD2a*, and *hopAW1a*). Instead, effectors in Pfm LV-5, namely *hopW1f* and *hopA1a*, which are able to suppress PTI, probably substitute for these effectors (Figure S5). A close ortholog of the positive control for the assay, AvrPtoB (HopAB1i from Pfm LV-5), may also be contributing to PTI-suppression but triggered an HR in *N. benthamiana* and thus its role could not be verified. Additionally, testing of Psa3 V-13 effectors HopD1a, AvrB2b, HopD2a, or HopAW1a, failed to complement pathogenicity in Pfm LV-5 (Figure S6). These results suggested instead that there might be effectors carried by Pfm LV- 5 that render it avirulent on ‘Hort16A’ plants.

The comparison of effector repertoires of Psa3 V-13 and Pfm LV-5 identified 16 effectors that are unique (absent in Psa3 V-13 or an allele present with <90% identity) to Pfm LV-5 with the potential to be avirulence effectors (Supplementary Table S3). Each of these 16 effectors was cloned under a synthetic *avrRps4* promoter with a C-terminal HA tag, and most validated for effector expression when delivered by Psa3 V-13 (Figure S7). Screening these Psa3 V-13 strains carrying Pfm LV-5 effectors on ‘Hort16A’ plants identified two effectors, *hopA1a* (10 fold reduction of pathogenicity) and *hopE1a* (100 fold reduction in pathogenicity) as candidate avirulence effectors (Figure S8) (Jayaraman *et al*., 2017). Delivering HopE1a_Pfm_ also largely eliminated Psa3 V-13-induced disease symptoms in ‘Hort16A’, while HopA1a_Pfm_ delivery barely reduced virulence (Figure S9). Since the C-terminal HA-tag may possibly interfere with immunity triggered by an effector, each of these effectors was cloned under their native promoters, where possible, and delivered by Psa3 V-13. Again only HopA1a_Pfm_ and HopE1a_Pfm_ were associated with a reduction of Psa3 V-13 pathogenicity in ‘Hort16A’ plants (Figure S10). Using a reporter eclipse assay for candidate avirulence effectors as well as effectors poorly expressed *in vitro*, also identified HopA1a_Pfm_ and HopE1a_Pfm_ as avirulence effectors in ‘Hort16a’ (Figure S11).

Knockout of avirulence effectors should allow for increased growth of Pfm LV-5 in ‘Hort16A’. Single and double knockout strains in Pfm LV-5 for *ΔhopA1a*, *ΔhopE1a*, or *ΔhopA1a*/*ΔhopE1a* were generated and tested for *in planta* growth. Surprisingly, none of these effector knockouts showed an increased pathogenicity in ‘Hort16A’ plants compared with wild-type Pfm LV-5 or Psa3 V-13 (Figure S12). Taken together, these findings suggest that Pfm LV-5 possesses all non-redundant virulence-associated effectors and its avirulence effectors do not contribute significantly to reduced pathogenicity in ‘Hort16A’ plants.

### Redundant pathogenicity-associated effectors from Psa3 largely target host RIN4 proteins

In an attempt to understand the lack of pathogenicity in Pfm LV-5 compared with Psa V-13, putatively redundant pathogenicity-associated effectors were tested for their contribution to host colonisation. Since HopA1a and HopE1a may be contributing to low rates of growth restriction, the double knockout strain Pfm LV-5 *ΔhopA1a*/*ΔhopE1a* was used for plasmid complementation of Psa3 V-13-specific effectors (*avrB2b*, *avrPto1b*, *avrRpm1a*, *hopD1a*, *hopF1c*, *hopH1a*, *hopZ5a*, *hopI1c*, *hopM1f*, *hopQ1a*, *hopF4a*, *hopBP1a*, *hopAM1a*, *hopD2a*, *hopAU1a*, *hopAW1a*, *hopF1e*, and *hopBN1a*). The expression of these HA-tagged Psa3 effectors in Pfm was validated under *hrp*-inducing conditions *in vitro* (Figure S13). Five Psa3 effectors were found to quantitatively increase pathogenicity of Pfm LV-5 *ΔhopA1a*/*ΔhopE1a* on ‘Hort16A’ plants (Figure 6A).

**Fig. 6.**
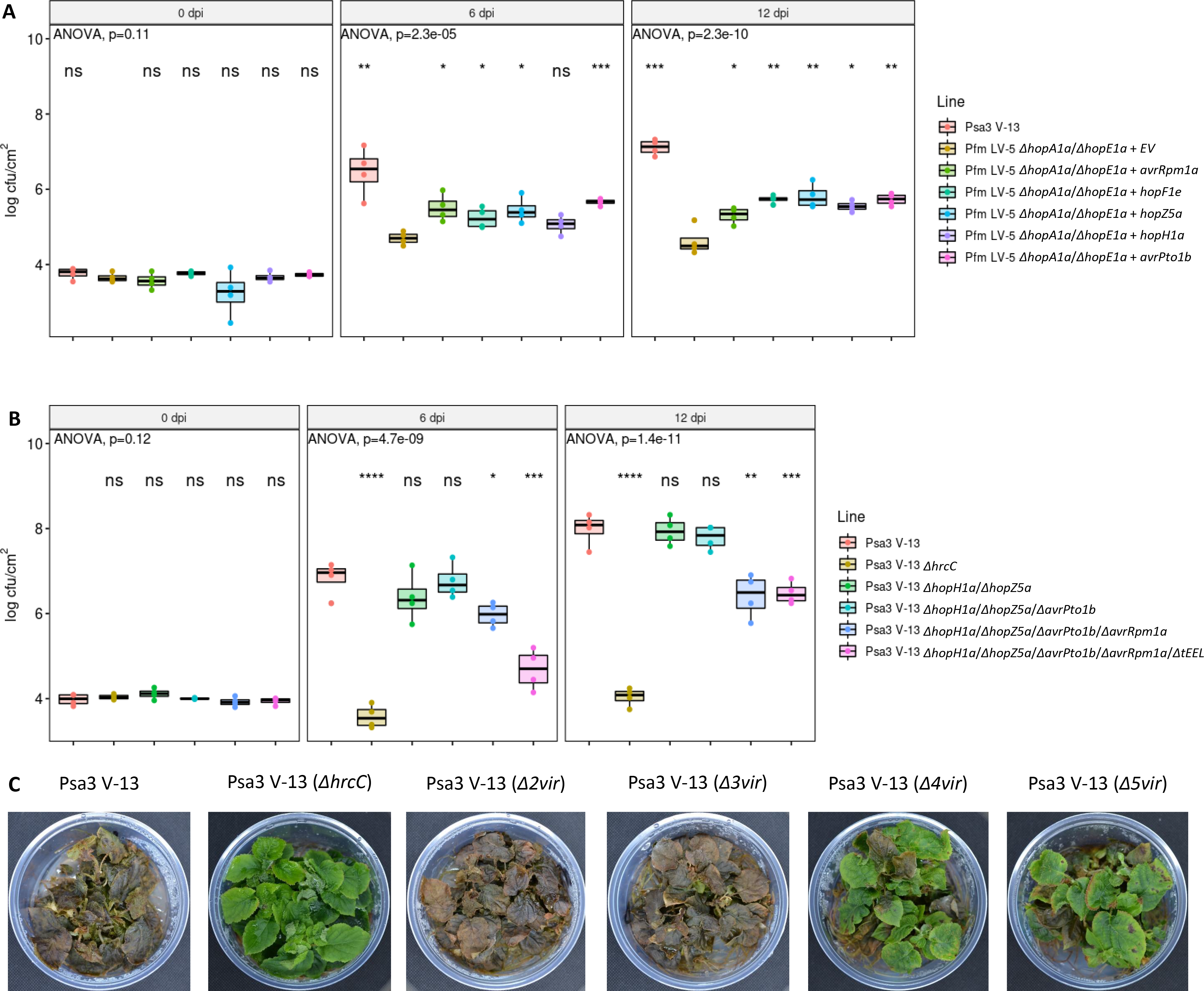
Five *Pseudomonas syringae* pv. *actinidiae* biovar 3 (Psa3) V-13 effectors offer redundant contributions to pathogenicity. **(A)** *Actinidia chinensis* var. *chinensis* ‘Hort16A’ plantlets were flood inoculated with wild-type Psa3 V-13, Pfm LV-5 Δ*hopA1a*/Δ*hopE1a* double mutant, or Pfm LV-5 Δ*hopA1a*/Δ*hopE1a* double mutant complemented with plasmid-borne Psa3 V-13 effectors *avrRpm1a*, *hopF1e*, *hopZ5a*, *hopH1a*, or *avrPto1b* at approximately 10 cfu/mL. Bacterial growth was quantified at 6 and 12 days post-inoculation by serial dilution and plate count quantification. Box and whisker plots, with black bars representing the median values and whiskers representing the 1.5 inter-quartile range, for *in planta* bacterial counts plotted as Log_10_ cfu/cm from four pseudobiological replicates. Asterisks indicate statistically significant differences from Welch’s t-test between the indicated strain and Pfm LV-5 Δ*hopA1*/Δ*hopE1* double mutant, where p≤.05 (*), p≤.01 (**), or p≤.001 (***); not significant (ns). These experiments were conducted three times on independent batches of ‘Hort16A’ plants, with similar results. **(B)** *Actinidia chinensis* var. *chinensis* ‘Hort16A’ plantlets were flood inoculated with wild-type Psa3 V-13, Δ*hrcC* mutant, Δ*hopH1a*/Δ*hopZ5a* mutant, Δ*hopH1a*/Δ*hopZ5a*/Δ*avrPto1b* mutant, Δ*hopH1a*/Δ*hopZ5a*/Δ*avrPto1b*/Δ*avrRpm1a* mutant, or Δ*hopH1a*/Δ*hopZ5a*/Δ*avrPto5a*/Δ*avrRpm1a*/Δ*tEEL* mutant at approximately 10 cfu/mL. Bacterial growth was quantified at 6 and 12 days post-inoculation by serial dilution and plate count quantification. Box and whisker plots, with black bars representing the median values and whiskers representing the 1.5 inter-quartile range, for *in planta* bacterial counts plotted as Log_10_ cfu/cm from four pseudobiological replicates. Asterisks indicate statistically significant differences from Welch’s t- test between the indicated strain and wild-type Psa3 V-13, where p≤.05 (*), p≤.01 (**), p≤.001 (***), or p≤.0001 (****); not significant (ns). These experiments were conducted three times on independent batches of ‘Hort16A’ plants, with similar results. **(C)** Symptom development on representative pottles of ‘Hort16A’ plantlets infected with strains in (B) at 50 days post-infection.

The five pathogenicity-associated Psa3 effectors (*hopZ5a*, *hopH1a*, *avrPto1b*, *avrRpm1a*, and *hopF1e*) that contribute to Pfm LV-5 *in planta* growth did not alter pathogenicity when knocked out individually in Psa3 V-13, suggesting that some redundancy across these effectors exists (Figure S14). To confirm this, cumulative knockouts of these effectors in Psa3 V-13 were generated and tested on ‘Hort16A’ plants. Notably, the Psa3 quadruple (*ΔhopH1a*/*ΔhopZ5a*/*ΔavrPto1b*/*ΔavrRpm1a*) and quintuple (*ΔhopH1a*/*ΔhopZ5a*/*ΔavrPto1b*/*ΔavrRpm1a*/*ΔtEEL*) knockouts showed a large drop in pathogenicity, confirming a redundant contribution of at least some of these effectors to pathogenicity (Figure 6B). The quadruple and quintuple mutants were also considerably reduced in virulence (Figure 6C).

Three out of these five putatively redundant effectors in Psa3, or their orthologs in other plant-pathogen systems, have recently been shown to target the plant immunity hub RIN4 (Yoon & Rikkerink, 2020; Choi *et al*., 2021; Jeleńska *et al*., 2021). HopF1e and HopH1 have not been characterised for their *in planta* targets, but HopF1e is part of the larger HopF family that has members known to interact with RIN4 (Lo *et al*., 2017). Indeed, Psa strains carry a large number of putative RIN4-targeting effectors (Hemara *et al*., 2022). These predicted Psa3 RIN4-targeting effectors, including HopF family effectors or Psa orthologs of known RIN4-targeting effectors that were not associated with increased Pfm LV-5 *ΔhopA1a*/*ΔhopE1a* growth, were tested for binding to RIN4 alleles from ‘Hort16A’. Co-immunoprecipitation screens were conducted in *N. benthamiana* plants with co-expression of alleles from one of three AcRIN4 loci and the effector of interest, as done previously for AvrRpm1a (Yoon & Rikkerink, 2020). The pathogenicity-associated effectors (HopZ5a, HopH1a, AvrPto1b, and HopF1e), known orthologs of RIN4 targeting effectors (HopBP1a and HopF1c), or putative HopF family effectors (HopF4a and HopBN1a) tested were all YFP-tagged and used to pull down any interacting FLAG-tagged RIN4 alleles (Figure 7). Notably, three out of the four redundant pathogenicity-associated effectors (HopZ5a, HopF1e, and AvrPto1b) pulled down at least two alleles of AcRIN4. No interaction was seen for HopH1a, similar to the negative control YFP alone. Meanwhile, both HopF1c and HopF4a also pulled down AcRIN4 alleles, but are not associated with increases in pathogenicity in Pfm LV-5, while HopBP1a and HopBN1a did not pull down AcRIN4 alleles. These results collectively suggest that AcRIN4 is a key target for a large number of Psa3 effectors that act together in a complex dynamic to facilitate ‘Hort16A’ infection.

**Fig. 7.**
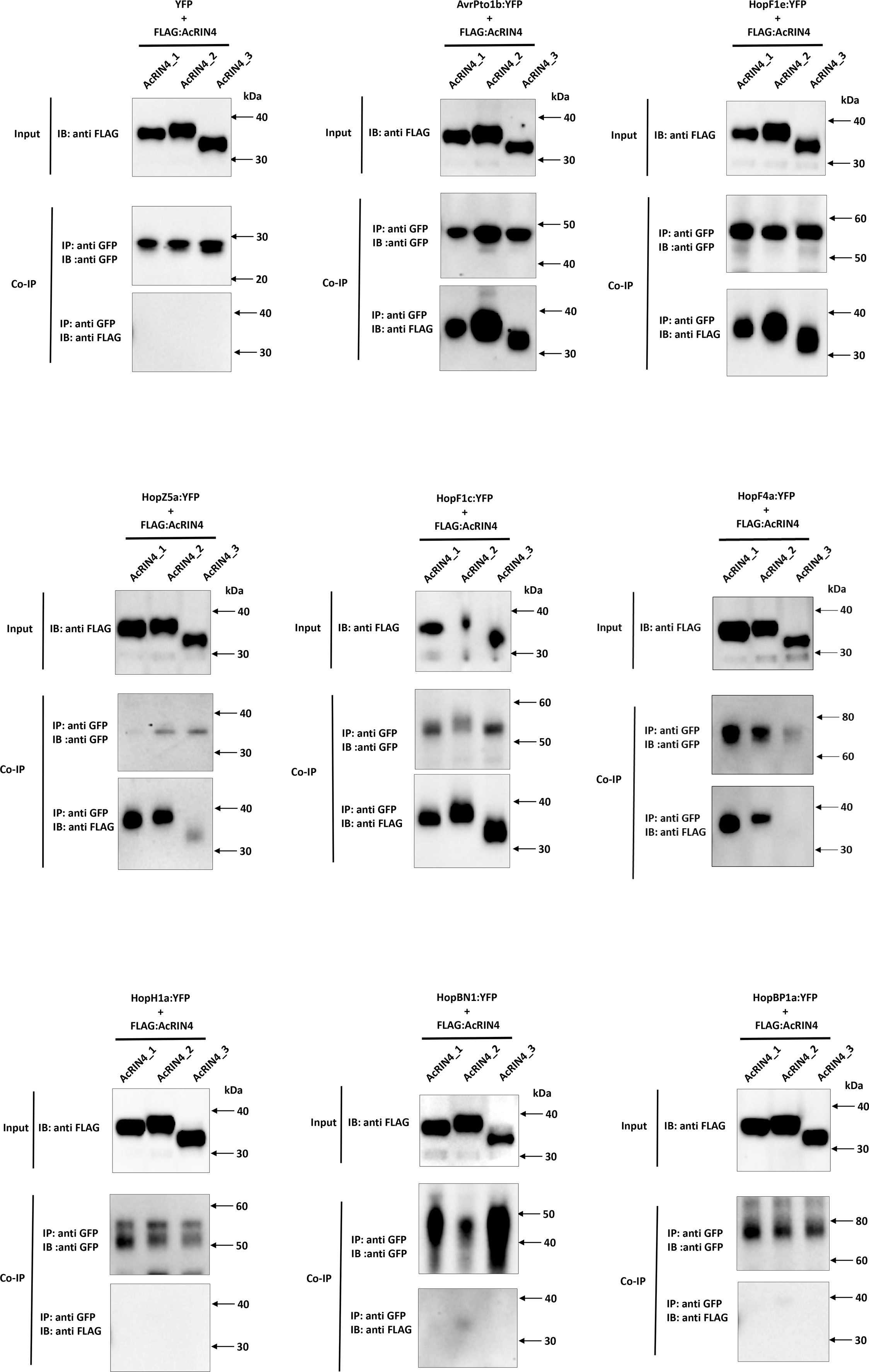
*Pseudomonas syringae* pv. *actinidiae* biovar 3 (Psa3) V-13 redundant virulence-associated effectors largely target kiwifruit RIN4 proteins. Co-immunoprecipitation of effectors AvrPto1b, HopF1e, HopZ5a, HopF1c, HopF4a, HopH1a, HopBN1a, or HopBP1a and target RIN4 proteins (AcRIN4- 1, -2, or -3). YFP-tagged effectors (or YFP alone) and FLAG-tagged RIN4 homologs were expressed simultaneously by *Agrobacterium tumefaciens*-mediated transient expression under a CaMV 35S promoter. Two days after infiltration, leaf samples were harvested, and protein extracts prepared and precipitated using anti-green fluorescent protein (GFP) antibody. A western blot from precipitated proteins was probed with anti-GFP (top) or anti-FLAG antibody (bottom). The plants were infiltrated with *Agrobacterium* at individual OD_600_ of 0.1. IP, co-immunoprecipitation; IB, immunoblotting.

## Discussion

This work sought to identify the virulence determinants that makes Psa3 strains hyper-virulent on the susceptible kiwifruit cultivar ‘Hort16A’, particularly in contrast to the various other *Pseudomonas* species that occupy the kiwifruit phyllosphere. Using a commensal, low virulence kiwifruit-colonising Pfm strain to search for virulence amplifiers in a double avirulence effector knockout strain (Pfm LV-5 Δ*hopA1a*/Δ*hopE1a)*, several redundantly acting effectors that largely target RIN4 were identified. While these effectors are collectively essential for full Psa3 virulence, no effectors were able to confer strong virulence to Pfm by themselves. This underscores the complexity of virulence in plant-colonising bacterial strains and suggests that factors beyond their effector repertoires may contribute to virulence.

For the well-characterised tomato and Arabidopsis pathogen Pto DC3000 in its infection of non-host *Nicotiana benthamiana*, several effectors were found to contribute towards pathogenicity following loss of avirulence effector HopQ1 (Kvitko *et al*., 2009; Cunnac *et al*., 2011; Wei *et al*., 2018). The AvrE1/HopM1/HopR1 redundant effector group (REG) was found to contribute to an aqueous apoplast and the AvrPto/AvrPtoB REG was found to target and suppress PTI (Kvitko *et al*., 2009). HopE1 supported increased growth *in planta*; HopG1 and HopAM1 were found to promote chlorotic and necrotic symptomology, respectively; and HopAA1 functioned redundantly with the phytotoxin coronatine to promote symptoms (Munkvold *et al*., 2009; Cunnac *et al*., 2011). The vast majority of Pto DC3000 effectors appear to have an ETI-suppression role (Jamir *et al*., 2004; Guo *et al*., 2009). In contrast, Psa3 appears to have little effector function in common with Pto DC3000. Notably, the contributions of AvrE1d and HopR1b that form a REG in Pto DC3000 have non-redundant roles in both pathogenicity and virulence on kiwifruit hosts (Jayaraman *et al*., 2020). In addition, this putative structure-related function is also seen in HopAS1b, which forms a similar potentially ‘pore-forming’ structure to AvrE1d/HopR1b, but appears not to be required for virulence or pathogenicity in kiwifruit plants (Figure 2; Figure S1). Notably, these three effectors were the only cell death-triggering Psa3 effectors that did not show a reduction in cell death upon silencing of *SGT1* in *N. benthamiana*, suggesting that all three are functional and that their virulence function may be associated with triggering cell death (Choi *et al*., 2017). Nevertheless, the variation among these three effectors’ requirements in Psa3 infection of kiwifruit plants suggests that the link between structural similarity and function is complex.

This work has identified two individual effectors (HopAZ1a and HopS2b) that contribute to pathogenicity (host colonisation) but have no effect on virulence (disease symptoms). This may be a unique role of these effectors that are not involved in symptom production, or a quantitative contribution that, despite affecting *in planta* accumulation, still allows for Psa3 colonisation beyond a threshold which then allows for symptom development (Stroud *et al*., 2022). Collectively, these non-redundant effectors or effector “sets” additively are essential for full pathogenicity and virulence in ‘Hort16A’ plants. Recently, HopAZ1a from Psa3 has been associated with targeting defence-associated PR5 and a cysteine peptidase Cp1 in kiwifruit plants (Zhu *et al*., 2022). This aforementioned work showed increased virulence for the *ΔhopAZ1a* mutant, unlike the reduced growth seen for our results.

However, their use of different cultivars of kiwifruit, and use of quantitative symptom development alone, may explain this discrepancy. Nevertheless, the finding that HopAZ1a targets secreted protein PR5 (and possibly Cp1) fits with previous work that showed HopAZ1a localised to what appears to be endoplasmic reticulum-like structures, suggesting that it may target defence-related secretion (Choi *et al*., 2017). While the plant targets of HopS2b are not known, a close ortholog of this effector from Pto DC3000 is a strong suppressor of ETI (Guo *et al*., 2009). Recently, the HopS family of ADP-ribosyl transferases were identified as significant contributors to virulence and ETI suppression through activity as NADases (Hulin & Ma, 2022). HopAZ1a and HopS2b appear to be the only effectors present universally across the five Psa biovars and Pfm, suggesting they play an important role in kiwifruit plant colonisation (McCann *et al*., 2013; Sawada & Fujikawa, 2019).

Using the same ‘single knockout’ approach used to identify the four non-redundant effectors contributing to pathogenicity of Psa3 on ‘Hort16A’, the exchangeable effector locus (EEL) was also identified as a significant contributor to pathogenicity, but not virulence. Interestingly, the entire extended EEL (xEEL) spanning ten effectors from *hopQ1a* to *hopF1a* was redundantly required for this contribution to pathogenicity. By using effector complementation, the xEEL was found to carry several functionally redundant effectors that participate in PTI-suppression: HopD1a, AvrB2b, HopAW1a, and HopD2a. These effectors appear to be able to facilitate PTI-suppression in the same way that AvrPto/AvrPtoB redundantly contribute to PTI suppression for Pto DC3000, and AvrPphB does for *P. syringae* pv. *phaseolicola* (Hann & Rathjen, 2007; Kvitko *et al*., 2009; Zhang *et al*., 2010).

Several redundant Psa3 effectors that could contribute to Pfm pathogenicity were identified: HopZ5a, AvrRpm1a, HopF1e, AvrPto5, and HopH1a. AvrRpm1a has been shown previously to target AcRIN4 alleles (Yoon & Rikkerink, 2020). In our current work, three out of the other four of these redundant virulence-associated effectors were shown to directly interact with AcRIN4 orthologs, suggesting a mechanism of virulence conserved in Psa3. This latter screen also tested all Psa3 effectors that are part of the HopF family (HopBN1a, HopF4a, HopF1c, and HopF1e), which has members known to target RIN4 as well as other orthologs that have been shown to target RIN4 (Wilton *et al*., 2010; Lo *et al*., 2017; Choi *et al*., 2021; Jeleńska *et al*., 2021). Several of these effectors showed interesting relationships between their ability to target AcRIN4 and their contribution to pathogenicity. HopH1a was the sole effector associated with virulence that did not bind to AcRIN4, while effectors HopF1c and HopF4a surprisingly did bind AcRIN4 but were not associated with pathogenicity (Figure 7). One reason that HopF1c was not identified is that it carries a defective SchF chaperone (Templeton *et al*., 2015). HopF1c has also previously been associated with triggering ETI in ‘Hort16A’ but is probably suppressed by other Psa3 effectors, so it is not surprising that it does not contribute to Pfm pathogenicity (Hemara *et al*., 2022). Meanwhile, the genomic sequence of *hopF4a* and its associated upstream *shcF* gene carries a transposon insertion that has deleted the ShcF N-terminus and separates *hopF4a* from its *hrpL* promoter (Templeton *et al*., 2015). This predicted disruption is corroborated by RNA-seq data showing HopF4a (previously named HopX3) does not appear to be expressed *in planta* (McAtee *et al*., 2018). HopF4a further lacks a functional catalytic triad from peptide sequence alignments and appears to be a non-functional member of the HopF family (Figure S15). HopBP1a (an ortholog of HopZ3 from *P. syringae* pv. *syringae* B728A that binds RIN4) did not bind to AcRIN4 and did not contribute to Pfm pathogenicity, albeit with undetectable expression when delivered by Pfm (Figure S13). Pfm LV-5 may require the AcRIN4-targeting capabilities supplied by these various effectors from Psa3 V-13 to increase Pfm pathogenicity in kiwifruit plants.

The ability of the RIN4-targeting set of effectors to increase Pfm pathogenicity implies they are also likely to be carrying out a similar role in Psa3. Why were these effectors not individually identified as contributing to pathogenicity or virulence in our screens? *A. arguta* plant lines like AA07_03 have evolved to recognise at least three of these effectors, and their deletion leads to an increase in fitness (Hemara *et al*., 2022). This illustrates the active role being played by these RIN4-interacting effectors in the evolution of Psa and kiwifruit germplasm near the likely point of origin of Psa as a species (McCann *et al*., 2017). In our analysis focussed on ‘Hort16A’, we suggest this is probably because of a complex series of interactions and active selection operating in both the host and pathogen around this important plant defence hub that is targeted by several different effector families across multiple bacterial plant pathogens (Sun *et al*., 2014; Rikkerink, 2018; Toruño *et al*., 2019). For example, in the case of the Psa3-susceptible ‘Hort16A’ host recognising hopF1c, the presence of this resistance (if now widespread in the wild kiwifruit-containing forests where Psa evolved) could well have resulted in selection for mutation of the associated chaperone ShcF to reduce effector delivery. Finally, evidence in this work and previously, suggests that there is a degree of redundancy among these effectors, as a cumulative loss of virulence on ‘Hort16A’ was evident in strains with multiple mutations in RIN4- interacting effectors (Figure 6; (Hemara *et al*., 2022)). Applying the principle of Occam’s razor would suggest their association with RIN4 is probably responsible for this redundancy.

A corollary question then becomes — what is the importance of targeting RIN4? Three effectors associated with RIN4 that trigger HR in *A. arguta* unusually did not trigger the ion leakage usually associated with this response (Hemara *et al*., 2022). This may indicate that one reason for targeting RIN4 is associated with suppressing ion leakage, a physiological response that is largely due to the loss of membrane integrity and is associated with the programmed cell death component of the HR. Thus it is conceivable that RIN4 performs an important regulatory function in controlling the initiation of (or limitation of) HR-associated cell death. The role of RIN4 in at least some hybrid-necrosis responses in lettuce could be an important functional clue here, albeit also probably in association with NLR proteins (Jeuken *et al*., 2009; Parra *et al*., 2016). The disordered protein structure of RIN4 has been suggested to be a key component that explains how it has evolved into such an important defence hub and target of post-translational modification by bacterial pathogens (Sun *et al*., 2014; Rikkerink, 2018). When our recent results are combined with previous research, it is increasingly clear that RIN4 is an equally import target for Psa (Yoon & Rikkerink, 2020).

## Acknowledgements

This work was funded by the Bio-protection Research Centre (Tertiary Education Commission) and the Royal Society Te Apārangi (including a Marsden FastStart grant to J.J.). We would like to thank Dr Jo Bowen (PFR) and Prof. Andrew Allan (PFR) for critical reading of this manuscript. The authors wish to acknowledge the use of New Zealand eScience Infrastructure (NeSI) high performance computing facilities as part of this research. New Zealand’s national facilities are provided by NeSI and funded jointly by NeSI’s collaborator institutions and through the Ministry of Business, Innovation & Employment’s Research Infrastructure programme. URL https://www.nesi.org.nz.

## Supplementary Materials

**Fig. S1.**
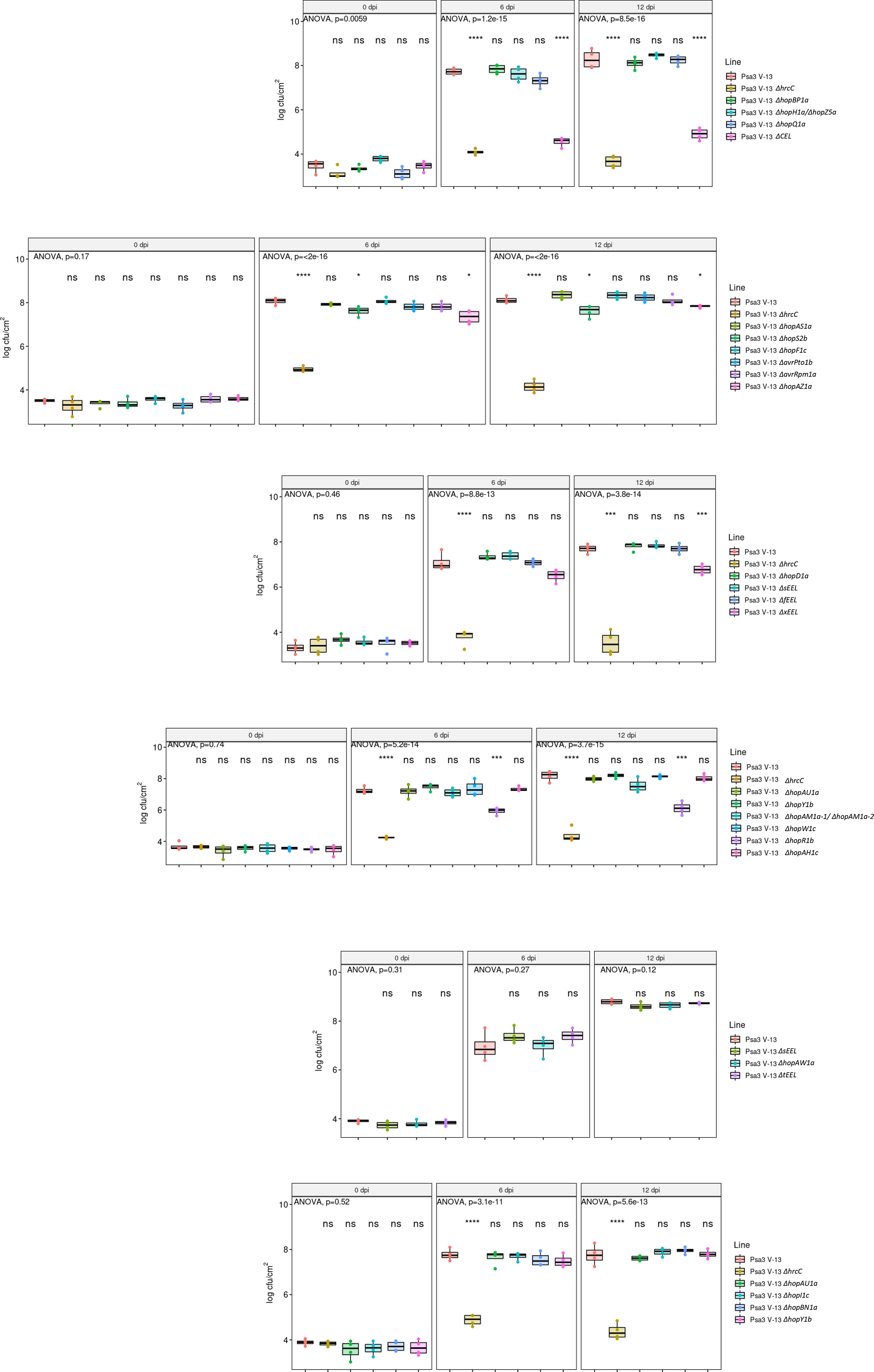
Screen of all single and block effector knockout strains. *Actinidia chinensis* var. *chinensis* ‘Hort16A’ plantlets were flood inoculated with wild-type Psa3 V-13, Δ*hrcC* mutant, or type III secreted effector mutants at approximately 10 cfu/mL. Bacterial growth was quantified at 6 and 12 days post-inoculation by serial dilution and plate count quantification. Box and whisker plots, with black bars representing the median values and whiskers representing the 1.5 inter-quartile range, for *in planta* bacterial counts plotted as Log_10_ cfu/cm^2^ from four pseudobiological replicates. Asterisks indicate statistically significant differences from Welch’s t-test between the indicated strain and wild-type Psa3 V-13, where p≤.05 (*), p≤.01 (**), p≤.001 (***). These experiments were conducted three times on independent batches of ‘Hort16A’ plants, with similar results.

**Fig. S2.**
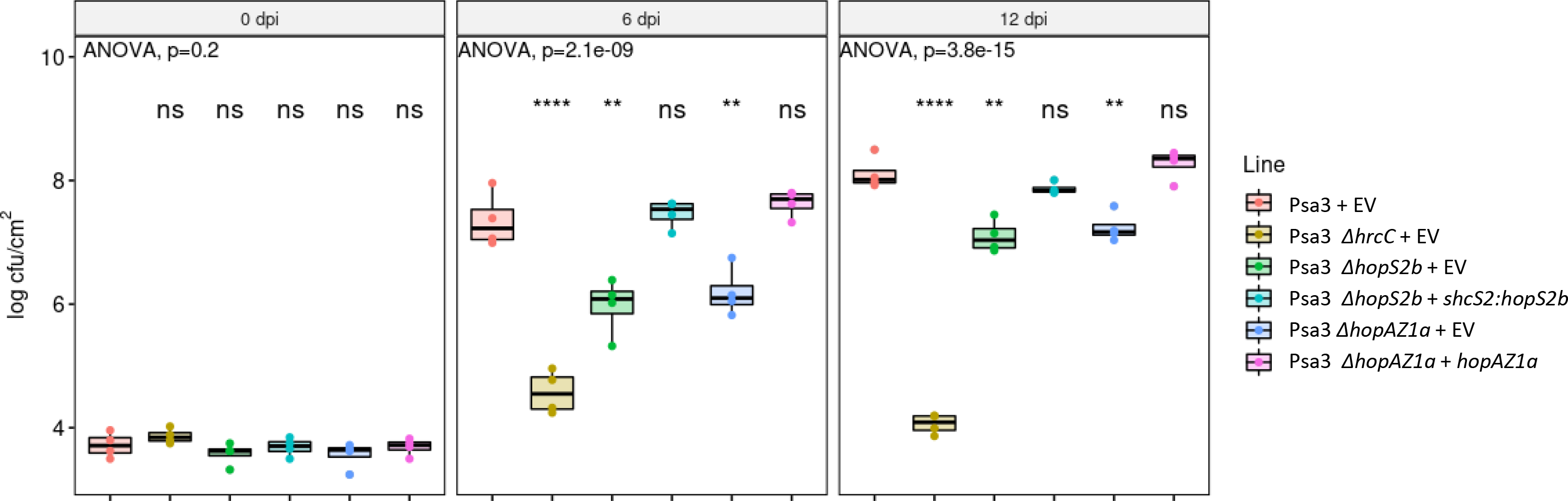
Plasmid complementation of *Pseudomonas syringae* pv. *actinidiae* biovar 3 (Psa3) V-13 effectors *hopS2b* and *hopAZ1a*. *Actinidia chinensis* var. *chinensis* ‘Hort16A’ plantlets were flood inoculated with wild-type Psa3 V-13 carrying an empty vector (+EV), Δ*hrcC* mutant +EV, Δ*hopS2b* mutant complemented with empty plasmid vector (+EV) or plasmid-borne *shcS2*:*hopS2b*, or Δ*hopAZ1* mutant complemented with empty plasmid vector (+EV) or plasmid-borne *hopAZ1a* at approximately 10 cfu/mL. Bacterial growth was quantified at 6 and 12 days post-inoculation by serial dilution and plate count quantification. Data are presented as box and whisker plots, with black bars representing the median values and whiskers representing the 1.5 inter-quartile range, for *in planta* bacterial counts plotted as Log_10_ cfu/cm^2^ from four pseudobiological replicates. Asterisks indicate statistically significant differences from Welch’s t-test between the indicated strain and Psa3 V-13 wild-type strain carrying empty vector, where p≤.01 (**), or p≤.0001 (****); not significant (ns). These experiments were conducted twice on independent batches of ‘Hort16A’ plants, with similar results.

**Fig. S3.**
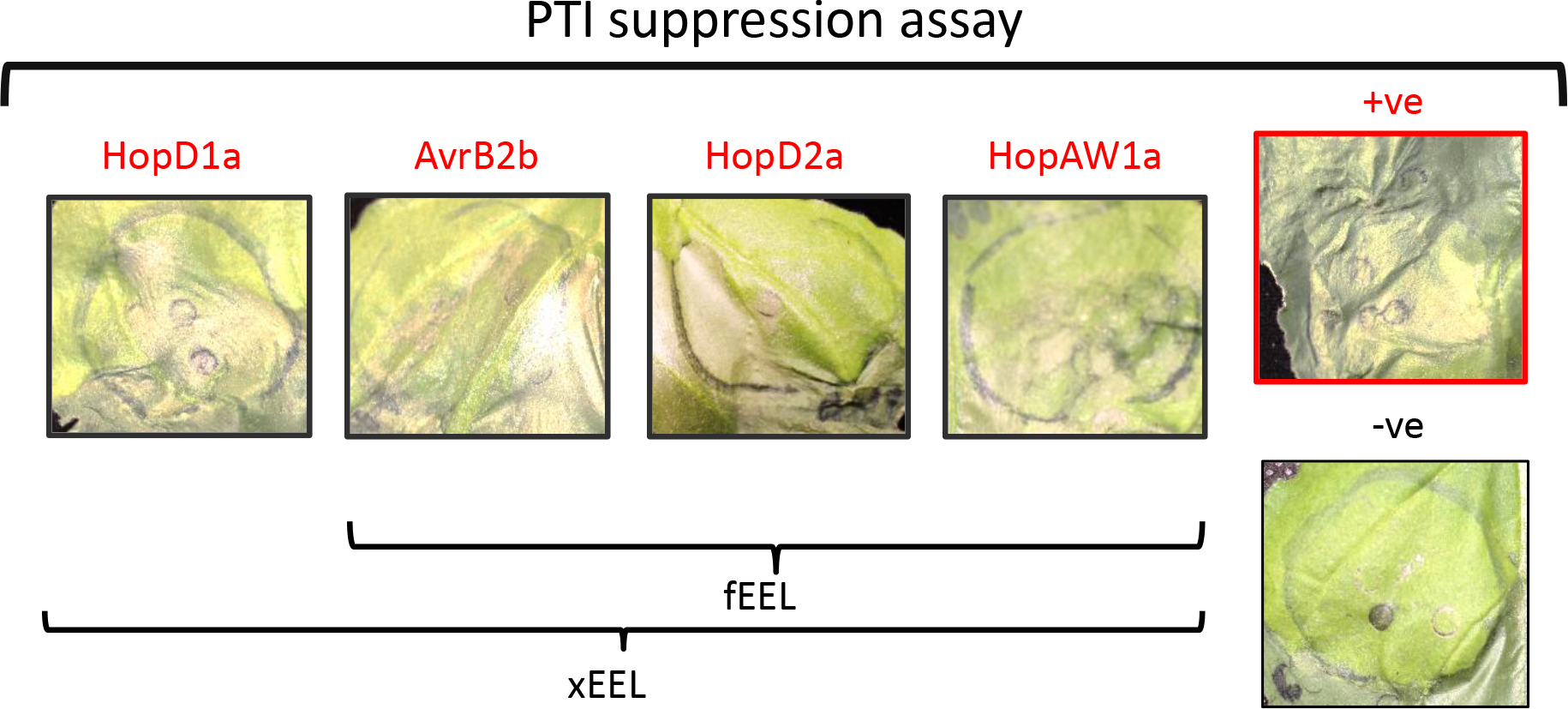
*Pseudomonas syringae* pv. *actinidiae* biovar 3 (Psa3) V-13 carries four PTI-suppressing effectors. The Psa3 V-13 effectors HopD1a, AvrB2b, HopD2a, and HopAW1a interfere with *P. fluorescens* (Pfo) Pf0-1(T3S)-induced PTI-mediated suppression of *P. syringae* pv. *tomato* (Pto) DC3000-triggered cell death. Leaves from 5-week-old *Nicotiana benthamiana* plants were infiltrated with Pfo Pf0-1(T3S) carrying *avrPtoB* (+ve), empty vector (-ve) or Psa3 V-13 effector (2 × 10^7^ cfu/mL) 8 h prior to *Pto* DC3000 (3 × 10^8^ cfu/mL) infection. Pto DC3000-triggered cell death was scored and photographed at 48 h post Pto DC3000 infection. This experiment was repeated three times, with similar results.

**Fig. S4.**
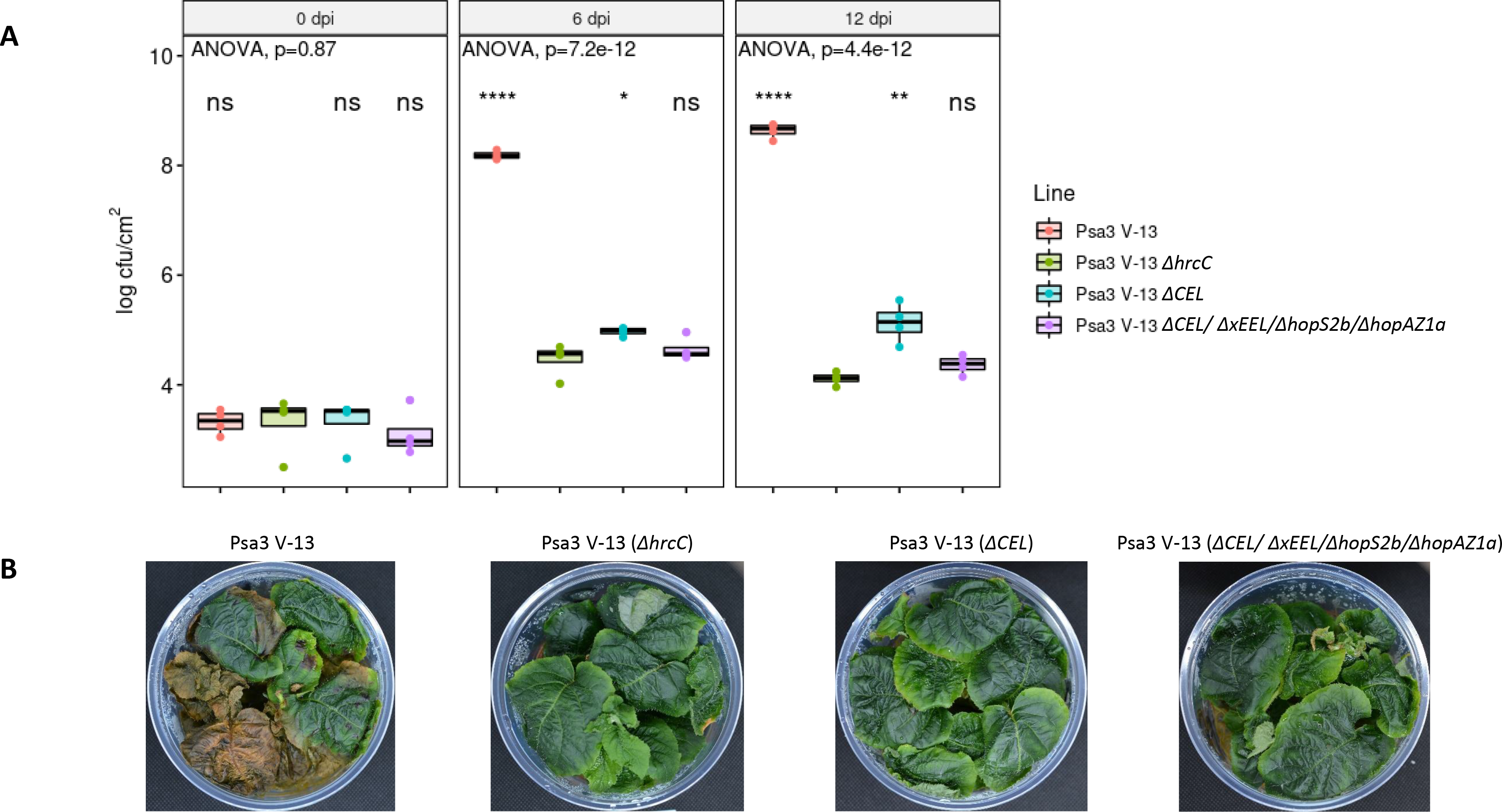
Loss of four *Pseudomonas syringae* pv. *actinidiae* biovar 3 (Psa3) V-13 pathogenicity-associated effector loci renders Psa3 V-13 non-pathogenic. *Actinidia chinensis* var. *chinensis* ‘Hort16A’ plantlets were flood inoculated with wild-type Psa3 V-13, Δ*hrcC* mutant, Δ*CEL* mutant, or Δ*CEL*/Δ*xEEL*/Δ*hopS2*/Δ*hopAZ1* quadruple mutant at approximately 10^6^ cfu/mL. (A) Bacterial growth was quantified at 6 and 12 days post-inoculation by serial dilution and plate count quantification. Box and whisker plots, with black bars representing the median values and whiskers representing the 1.5 inter-quartile range, for *in planta* bacterial counts plotted as Log_10_ cfu/cm^2^ from four pseudobiological replicates. Asterisks indicate statistically significant differences from Welch’s t-test between the indicated strain and the Psa3 V-13 Δ*hrcC* mutant, where p≤.05 (*), p≤.01 (**), or p≤.0001 (****); not significant (ns). (B) Symptom development on representative pottles of ‘Hort16A’ plantlets infected with strains in (A) at 50 days post-infection.

**Fig. S5.**
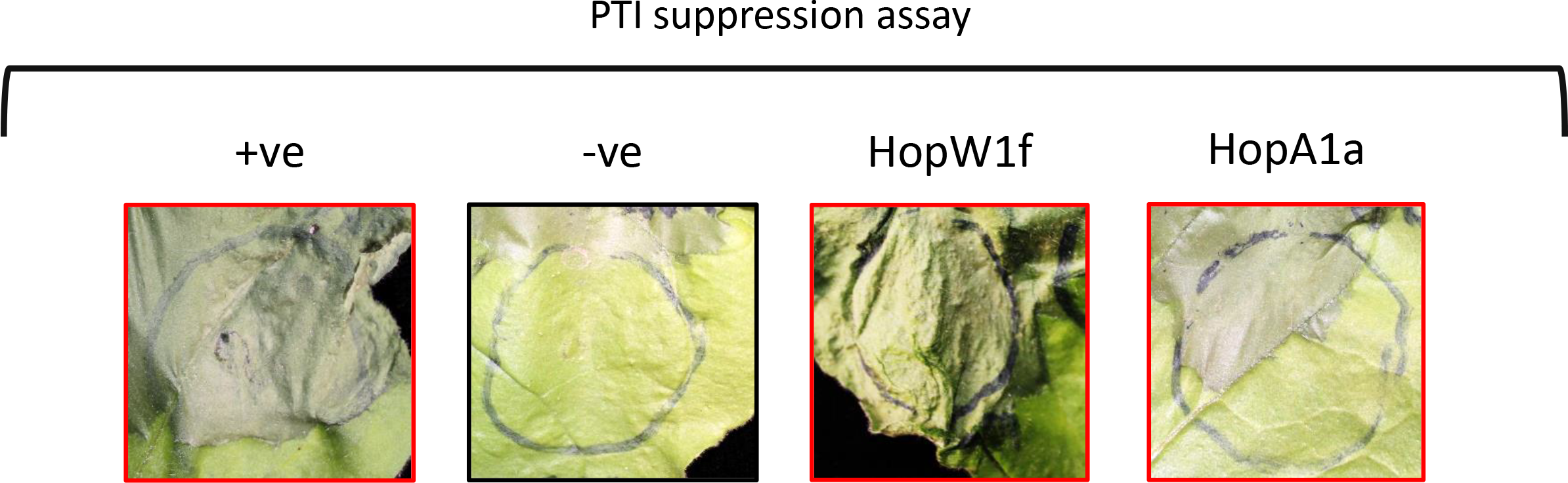
At least two PTI-suppressing effectors are carried by *Pseudomonas syringae* pv. *actinidifoliorum* (Pfm) LV-5. The Pfm LV-5 effectors HopW1f and HopA1a interfere with *P. fluorescens* (Pfo) Pf0-1(T3S)-induced PTI-mediated suppression of *P. syringae* pv. *tomato* (Pto) DC3000-triggered cell death. Leaves from 5-week-old *Nicotiana benthamiana* plants were infiltrated with Pfo Pf0-1(T3S) carrying *avrPtoB* (+ve), empty vector (-ve) or Pfm LV-5 effector (2 × 10 cfu/mL) 8 h prior to *Pto* DC3000 (3 × 10 cfu/mL) infection. Pto DC3000-triggered cell death was scored and photographed at 48 h post Pto DC3000 infection. This experiment was repeated twice, with similar results. HopAB1i could not be tested since it triggers a strong cell death response in *N. benthamiana* plants.

**Fig. S6.**
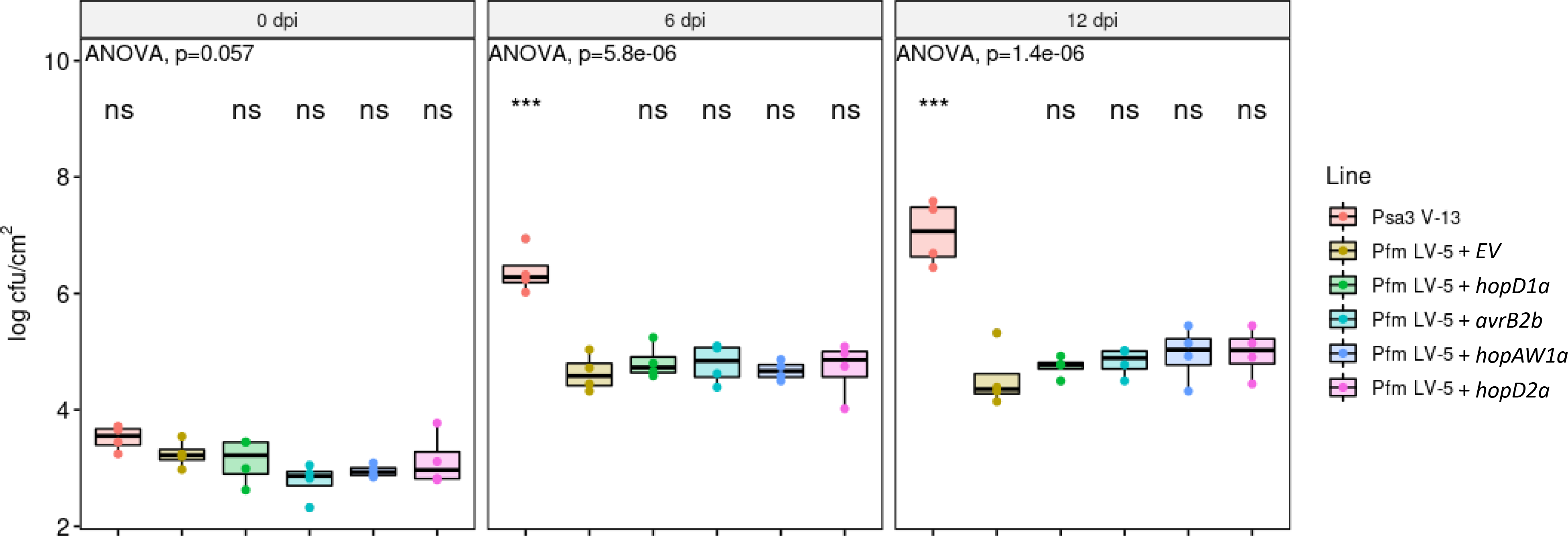
The four *Pseudomonas syringae* pv. *actinidiae* biovar 3 (Psa3) V-13 PTI-suppressing extended exchangeable effector locus (xEEL) effectors are unable to complement *P. syringae* pv. *actinidifoliorum* LV-5 pathogenicity. *Actinidia chinensis* var. *chinensis* ‘Hort16A’ plantlets were flood inoculated with wild-type Psa3 V-13 carrying empty vector (+ EV), Pfm LV-5 +EV, or Pfm LV-5 carrying plasmid-borne Psa3 V-13 effectors *hopD1a*, *avrB2b*, *hopAW1a*, or *hopD2a* at approximately 10 cfu/mL. Bacterial growth was quantified at 6 and 12 days post-inoculation by serial dilution and plate count quantification. Box and whisker plots, with black bars representing the median values and whiskers representing the 1.5 inter-quartile range, for *in planta* bacterial counts plotted as Log_10_ cfu/cm^2^ from four pseudobiological replicates. Asterisks indicate statistically significant differences from Welch’s t-test between the indicated strain and Pfm LV-5 strain carrying empty vector, where p≤.001 (***); not significant (ns).

**Fig. S7.**
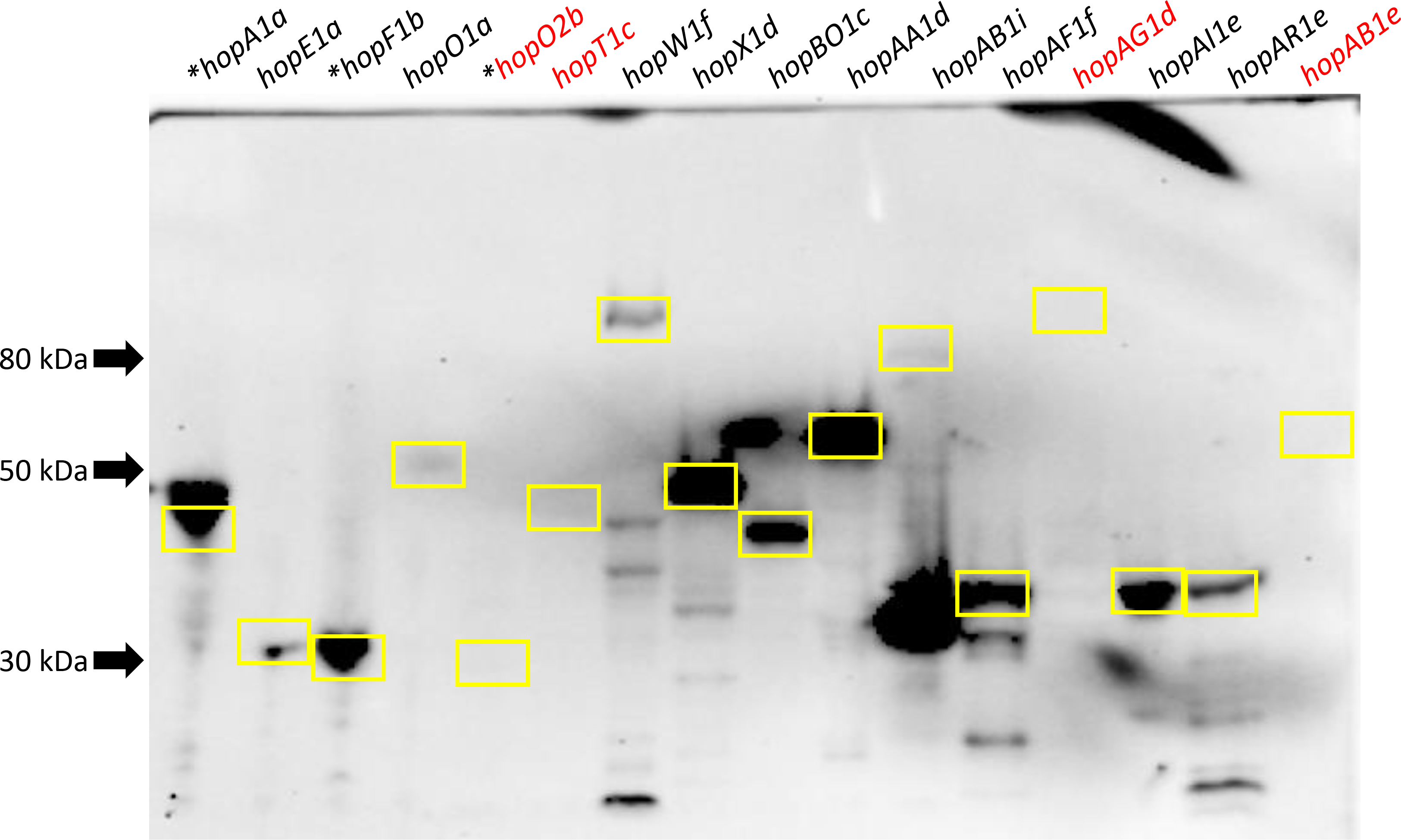
Secretion of *Pseudomonas syringae* pv. *actinidifoliorum* (Pfm) LV-5 effectors by plasmid complementation in *P. syringae* pv. *actinidiae* biovar 3 (Psa3) V-13 during expression *in vitro*. Psa3 V-13 carrying type III secreted effector proteins tagged with 6 × HA were diluted to 5 x 10 cfu/mL in *hrp*-inducing liquid medium, samples harvested at 6 h post-inoculation by centrifugation at 12,000 *g*, boiled in 1x Laemmli buffer, and western blots conducted using α-HA antibody. Yellow boxes indicate expected sizes for each tagged protein band. Asterisks indicate the effector is cloned with its preceding chaperone. Red font for effector label indicates non-detectable/weak band at the expected size.

**Fig. S8.**
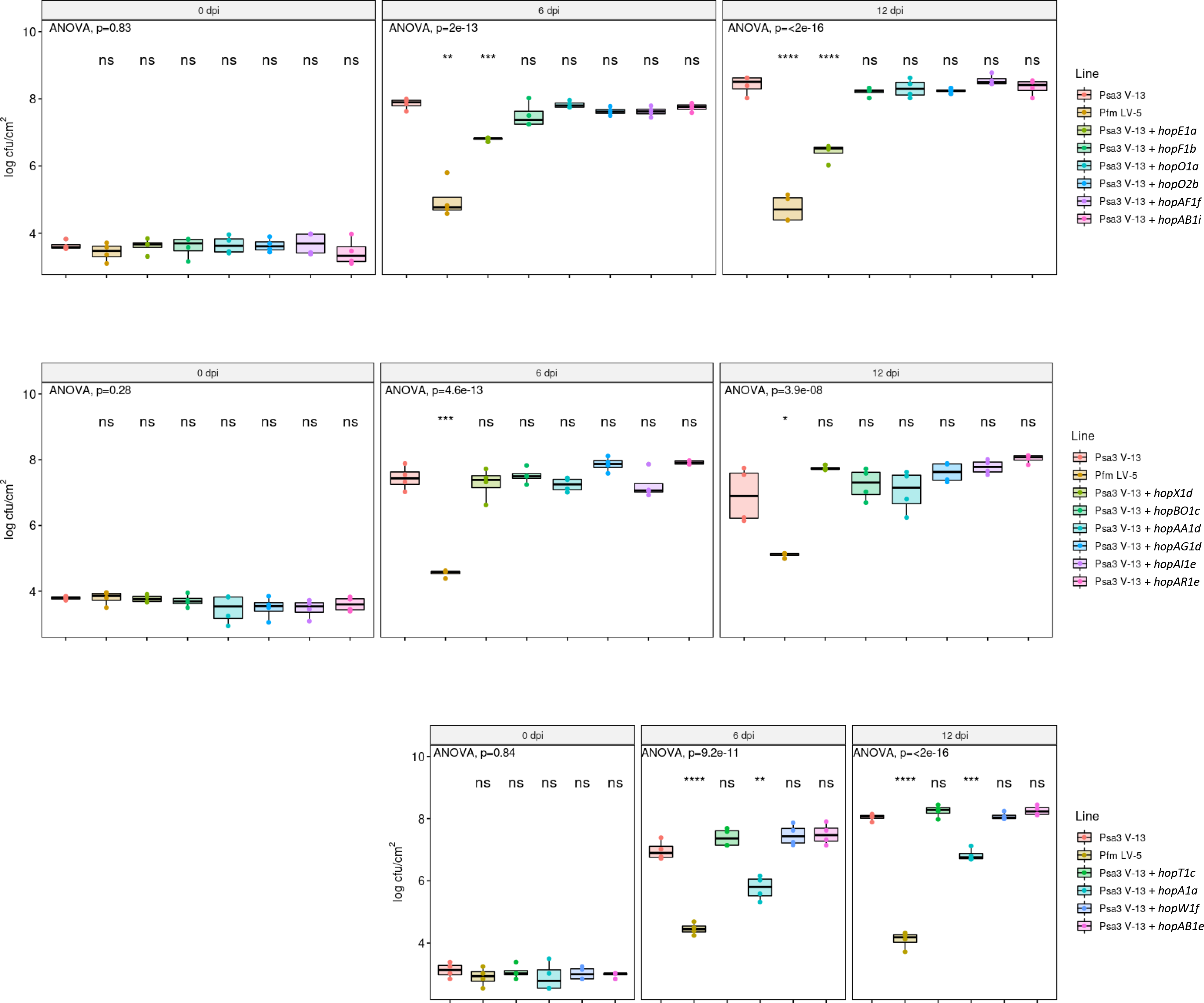
Screen of all *Pseudomonas syringae* pv. *actinidifoliorum* (Pfm) LV-5 unique effectors in *P. syringae* pv. *actinidiae* biovar 3 (Psa3) V-13. *Actinidia chinensis* var. *chinensis* ‘Hort16A’ plantlets were flood inoculated with wild-type Psa3 V-13, Pfm LV-5, or Psa3 V-13 complemented with a plasmid-borne type III secreted effector unique to Pfm LV-5 cloned under a synthetic *avrRps4* promoter and tagged with a 6xHA tag at approximately 10 cfu/mL. Bacterial growth was quantified at 6 and 12 days post-inoculation by serial dilution and plate count quantification. Box and whisker plots, with black bars representing the median values and whiskers representing the 1.5 inter-quartile range, for *in planta* bacterial counts plotted as Log_10_ cfu/cm^2^ from four pseudobiological replicates. Asterisks indicate statistically significant differences from Welch’s t-test between the indicated strain and wild-type Psa3 V-13, where p≤.01 (**), p≤.001 (***), p≤.0001 (****). These experiments were conducted twice on independent batches of ‘Hort16A’ plants, with similar results.

**Fig. S9.**
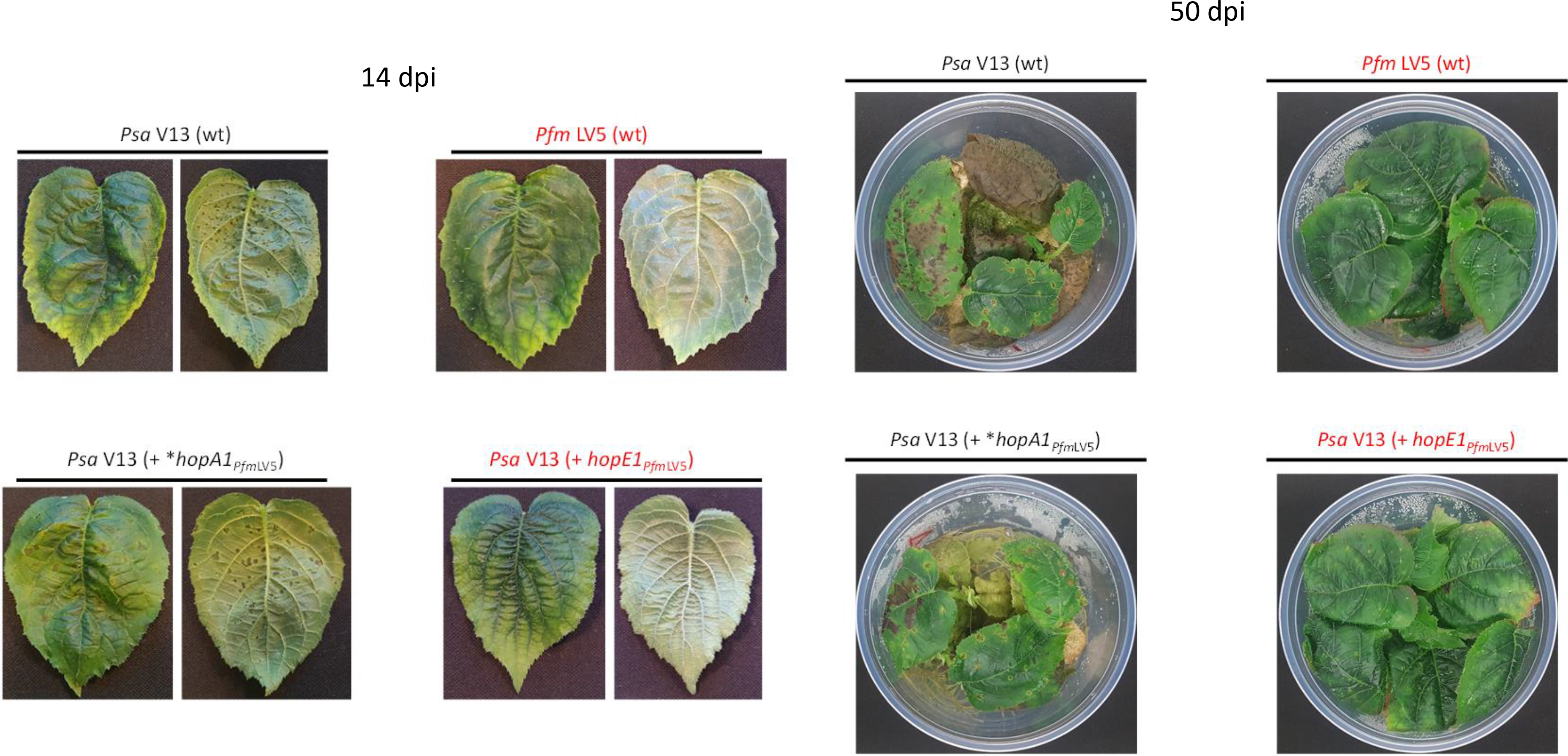
*Pseudomonas syringae* pv. *actinidiae* biovar 3 (Psa3) V-13 complemented with *P. syringae* pv. *actinidifoliorum* (Pfm) LV-5 avirulence effector *hopE1a* but not *hopA1a* lacks virulence *in planta*. *Actinidia chinensis* var. *chinensis* ‘Hort16A’ plantlets were flood inoculated with wild-type Psa3 V-13 (wt), Pfm LV-5 (wt), Psa3 V-13 carrying plasmid-borne *hopE1a* (*avrRps4* promoter, 6xHA tagged), or Psa3 V-13 carrying plasmid-borne *hopA1a* (*avrRps4* promoter, 6xHA tagged; asterisk indicates cloning of whole operon including *shcA*) at approximately 10 cfu/mL. Photographs of symptom development on representative pottles of ‘Hort16A’ plantlets were taken at 14 days post-infection (dpi) for individual leaves both abaxially and adaxially, and of full pottles at 50 dpi. Leaves and pottles that show lack of symptom development are indicated with red labels.

**Fig. S10.**
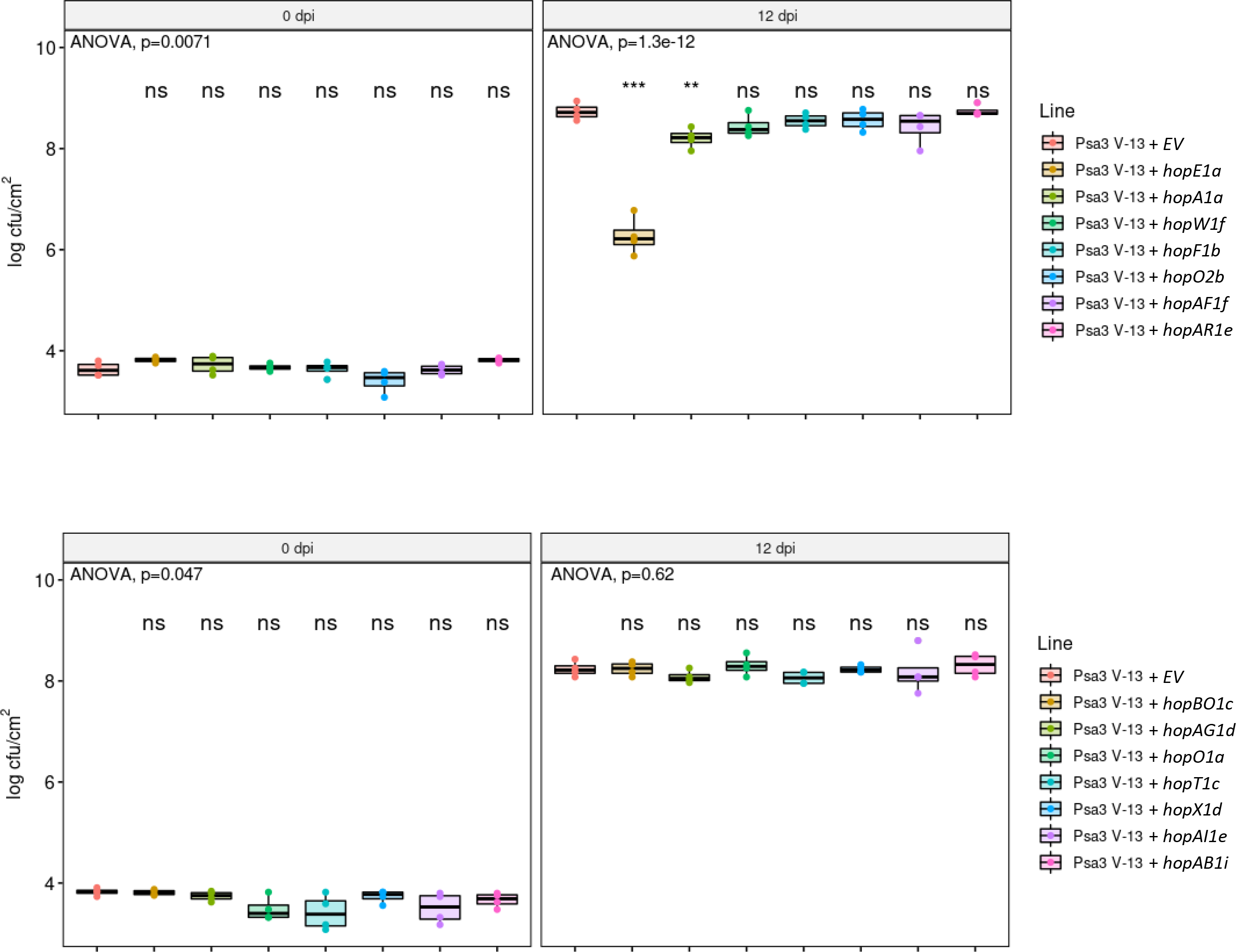
Screen of *Pseudomonas syringae* pv. *actinidifoliorum* (Pfm) LV-5 unique effectors in *P. syringae* pv. *actinidiae* biovar 3 (Psa3) V-13. *Actinidia chinensis* var. *chinensis* ‘Hort16A’ plantlets were flood inoculated with wild-type Psa3 V-13 complemented with an empty vector (EV) or Psa3 V-13 complemented with a plasmid-borne type III secreted effector unique to Pfm LV-5 cloned under its native promoter (with no tag) at approximately 10 cfu/mL. Bacterial growth was quantified at 12 days post-inoculation by serial dilution and plate count quantification. Box and whisker plots, with black bars representing the median values and whiskers representing the 1.5 inter-quartile range, for *in planta* bacterial counts plotted as Log_10_ cfu/cm^2^ from four pseudobiological replicates. Asterisks indicate statistically significant differences from Welch’s t-test between the indicated strain and wild-type Psa3 V-13, where p≤.01 (**), or p≤.001 (***); not significant (ns). Effectors *hopO1a*, *hopT1c*, *hopX1d*, and *hopAI1e* are cloned under a synthetic *avrRps4* promoter since their native promoter could not be cloned because of their presence in an operon. Effectors *hopAA1d* and *hopAB1e* could not be cloned under their native promoters.

**Fig. S11.**
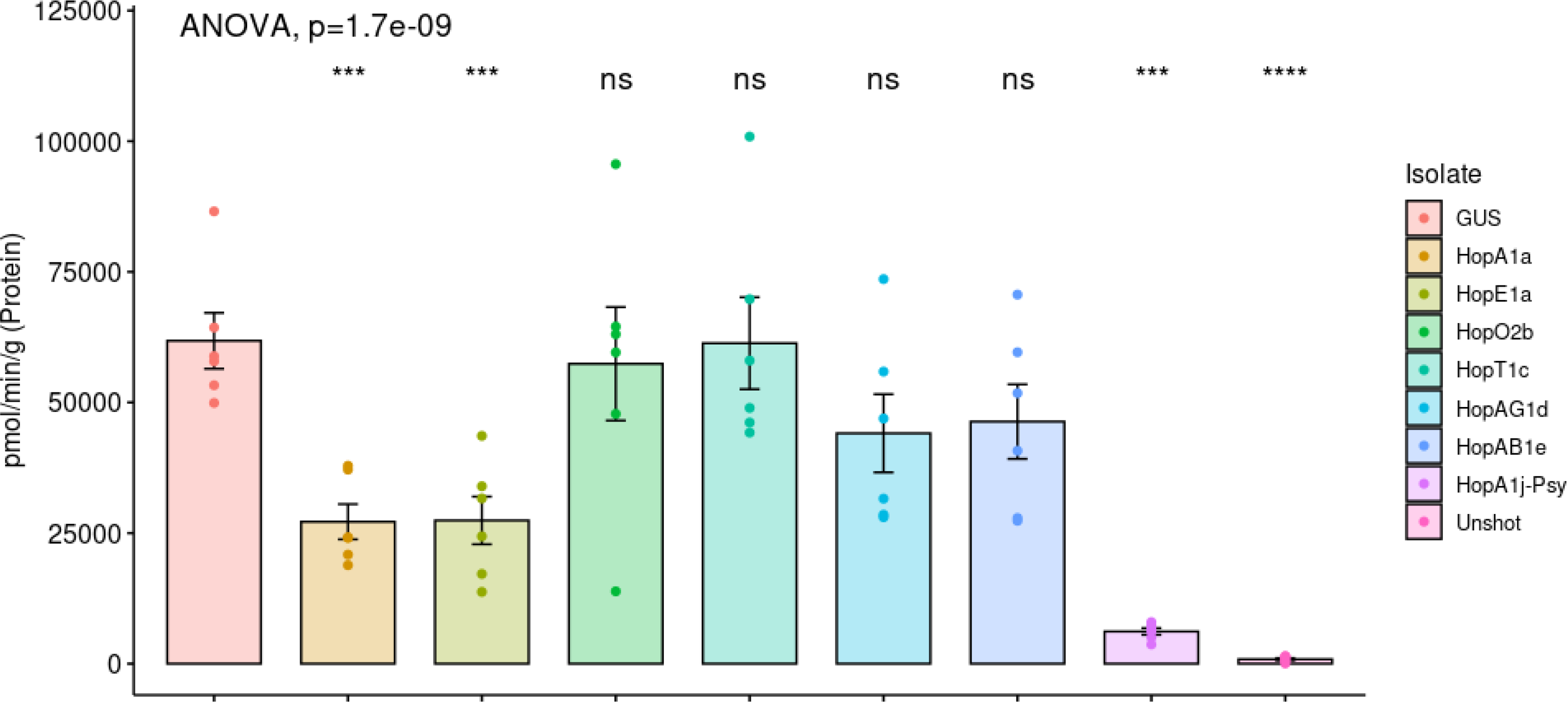
Biolistic transformation reporter eclipse assays for *Pseudomonas syringae* pv. *actinidifoliorum* (Pfm) LV-5 effectors demonstrate hypersensitive response triggered by HopE1a. Putative Pfm LV-5 avirulence effector genes (*hopA1a* or *hopE1a*) or effector genes with unclear *Pseudomonas* expression (*hopO2b*, *hopT1c*, *hopAB1e*, *hopAG1d*) cloned on binary vector constructs tagged with YFP, or an empty vector (labelled as GUS), were co-expressed with a β-glucuronidase (GUS) reporter construct using biolistic bombardment and priming in leaves from *Actinidia chinensis* var. *chinensis* ‘Hort16A’ plantlets. The GUS activity was measured 48 h after DNA bombardment. Error bars represent the standard errors of the means for six technical replicates each (n=6). *hopA1j* cloned similarly from *P. syringae* pv. *syringae* 61 was used as positive control and un-infiltrated leaf tissue (Unshot) as negative control. Statistical significance is indicated for a one-way ANOVA and Tukey’s HSD *post hoc* test : p≤.001 (***), p≤.0001 (****), and p>.05 (ns).

**Fig. S12.**
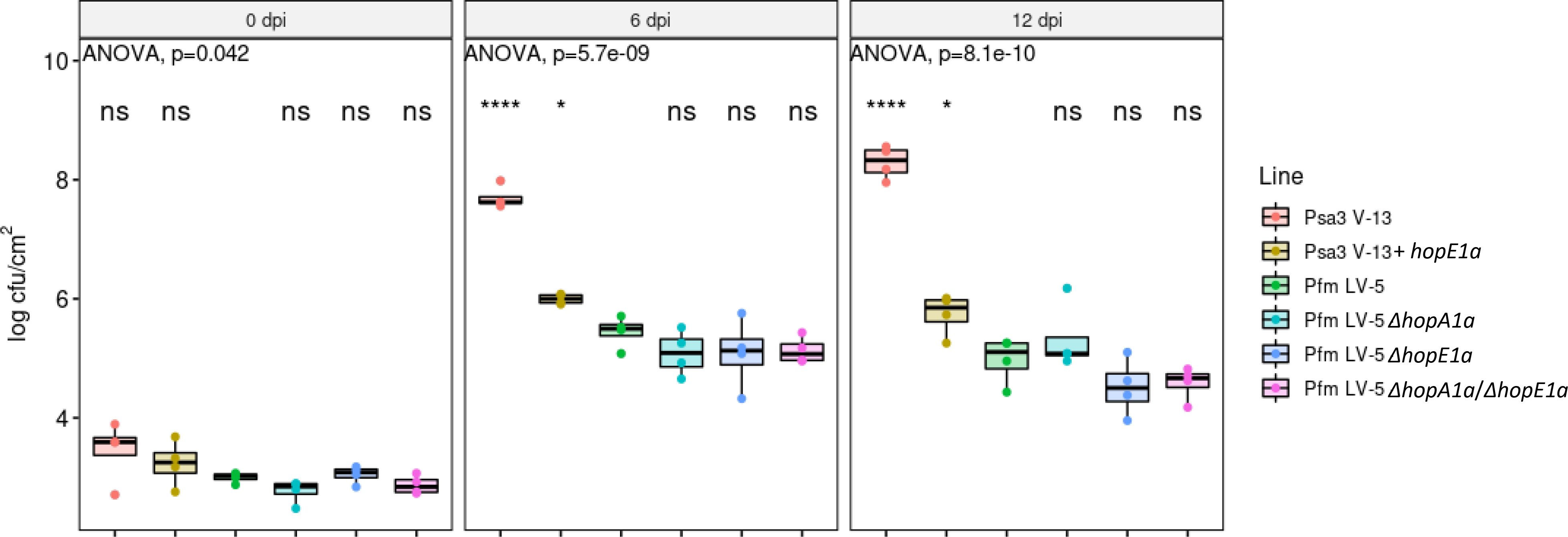
Deletion of both *Pseudomonas syringae* pv. *actinidifoliorum* (Pfm) LV-5 avirulence effectors does not confer increased pathogenicity. *Actinidia chinensis* var. *chinensis* ‘Hort16A’ plantlets were flood inoculated with wild-type Psa3 V-13, Psa3 V-13 carrying avirulence effector *hopE1a*, wild-type Pfm LV-5, Pfm LV-5 Δ*hopA1* mutant, Pfm LV-5 Δ*hopE1* mutant, or Pfm LV-5 Δ*hopA1*/Δ*hopE1* double mutant at approximately 10 cfu/mL. Bacterial growth was quantified at 6 and 12 days post-inoculation by serial dilution and plate count quantification. Box and whisker plots, with black bars representing the median values and whiskers representing the 1.5 inter-quartile range, for *in planta* bacterial counts plotted as Log_10_ cfu/cm^2^ from four pseudobiological replicates. Asterisks indicate statistically significant differences from Welch’s t-test between the indicated strain and the Pfm LV-5 strain, where p≤.05 (*) or p≤.0001 (****); not significant (ns).

**Fig. S13.**
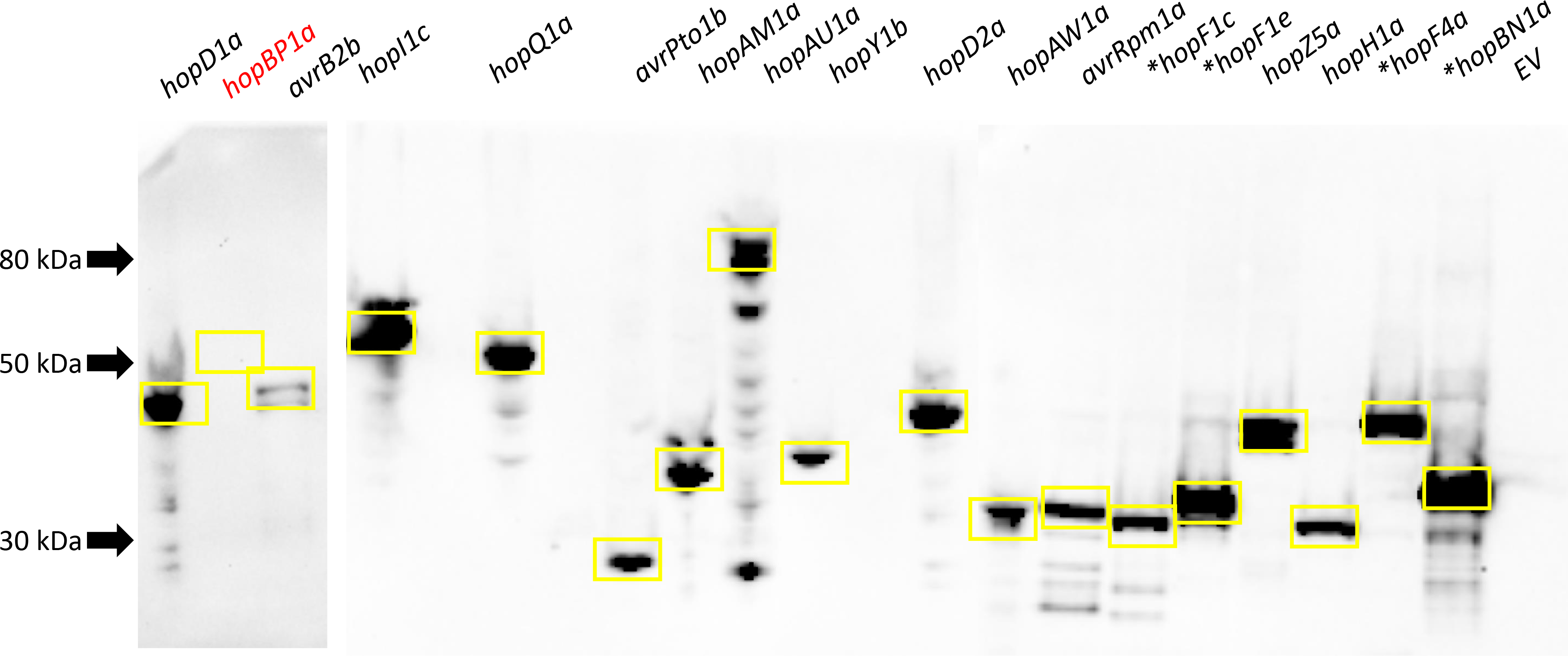
Secretion of *Pseudomonas syringae* pv. *actinidiae* biovar 3 (Psa3) V-13 effectors by plasmid complementation in *P. syringae* pv. *actinidifoliorum* (Pfm) LV-5 during expression *in vitro*. Pfm LV-5 carrying type III secreted effector proteins tagged with 6 × HA were diluted to 5 x 10^8^ cfu/mL in *hrp*- inducing liquid medium, samples harvested at 6 h post-inoculation by centrifugation at 12,000 *g*, boiled in 1x Laemmli buffer, and western blots conducted using α-HA antibody. Yellow boxes indicate expected sizes for each tagged protein band. Asterisks indicate the effector is cloned with its preceding chaperone. Red font for effector label indicates non-detectable/weak band at the expected size.

**Fig. S14.**
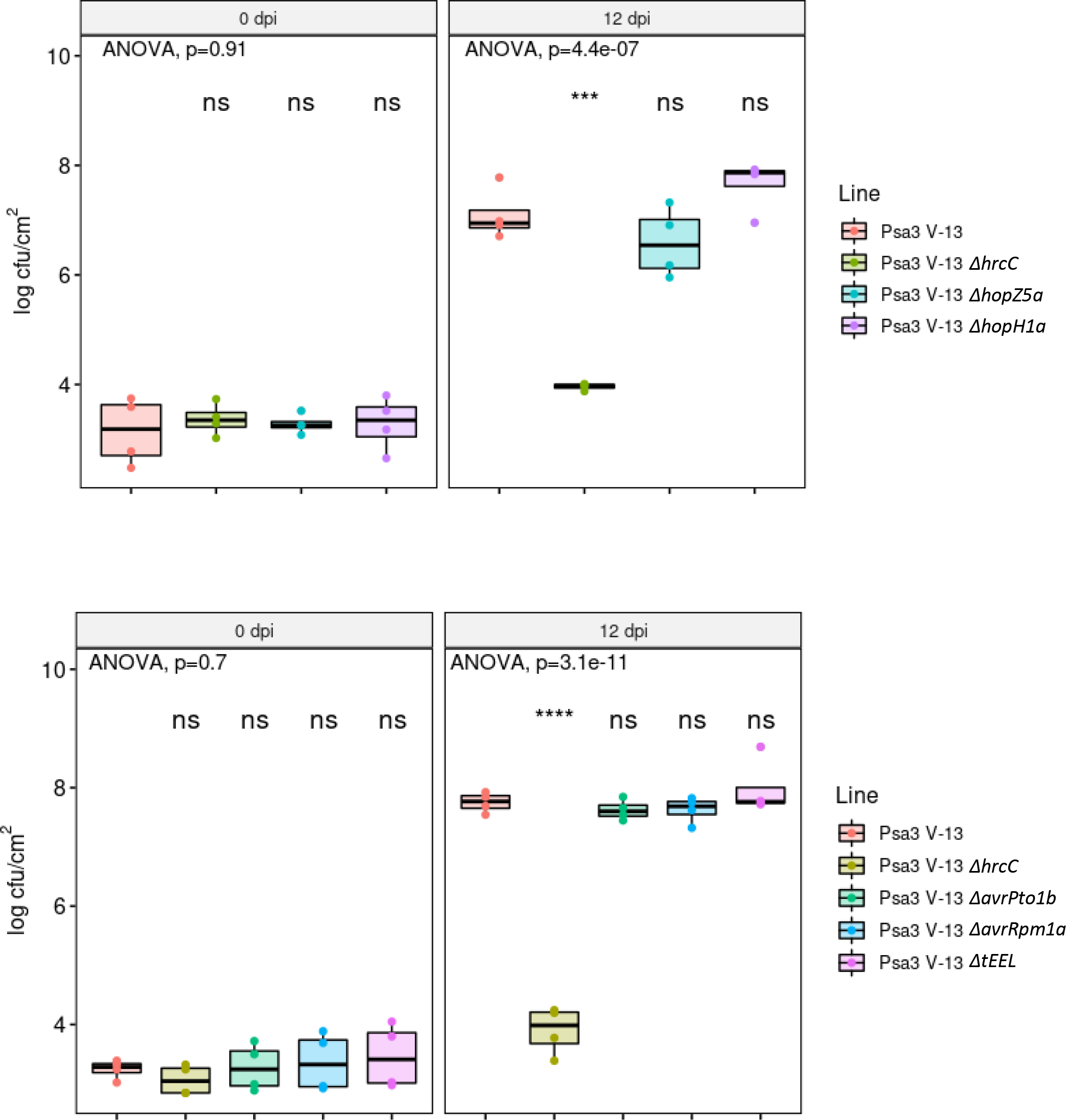
Separate effector knockouts in *Pseudomonas syringae* pv. *actinidiae* biovar 3 (Psa3) V-13 for effectors that complement pathogenicity in *P. syringae* pv. *actinidifoliorum* (Pfm) LV-5 demonstrates effector redundancy. *Actinidia chinensis* var. *chinensis* ‘Hort16A’ plantlets were flood inoculated with wild-type Psa3 V-13, Δ*hrcC* mutant, or indicated type III secreted effector mutants at approximately 10 cfu/mL. The Δ*tEEL* mutant spans *hopF1e*, *hopAF1b, hopD2a*, and *hopF1a*. Bacterial growth was quantified at 12 days post-inoculation by serial dilution and plate count quantification. Box and whisker plots, with black bars representing the median values and whiskers representing the 1.5 inter-quartile range, for *in planta* bacterial counts plotted as Log_10_ cfu/cm^2^ from four pseudobiological replicates. Asterisks indicate statistically significant differences from Welch’s t-test between the indicated strain and wild-type Psa3 V-13, where p≤.001 (***), or p≤.0001 (****); not significant (ns).

**Fig. S15.**
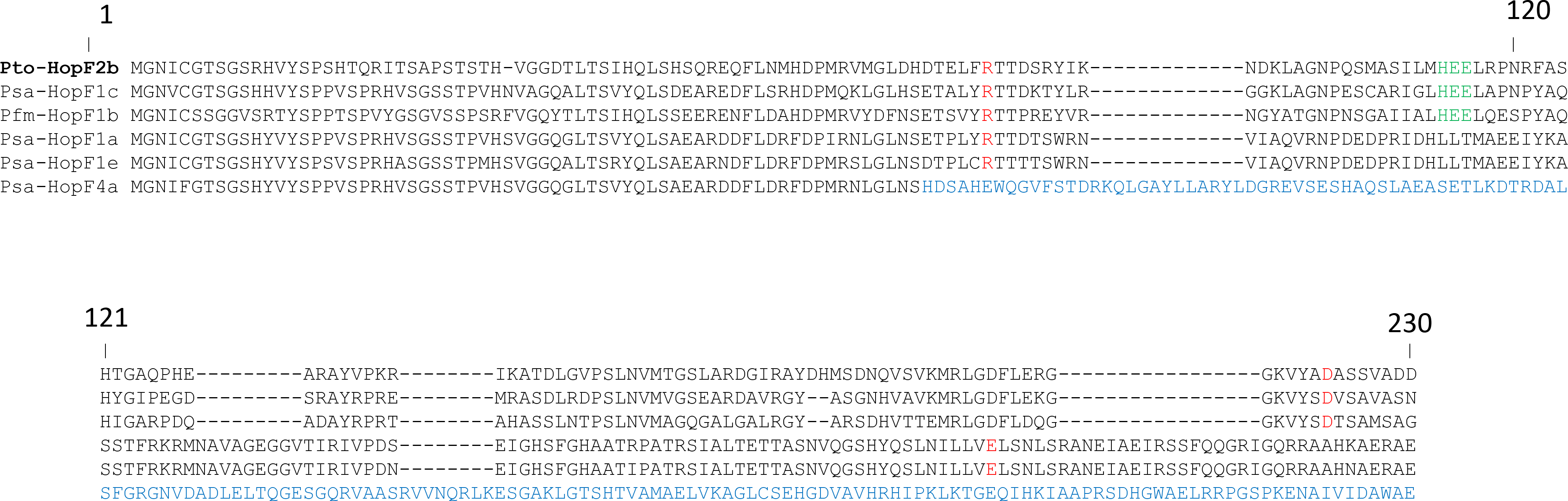
Alignment of HopF family effectors from Pto, Psa and Pfm identify HopF4a as a non-functional effector. Amino acid sequence alignment of HopF2 (HopF2b) from *P. syringae* pv. *tomato* [Pto] DC3000 (set as reference, bold); HopF1c, HopF1a, HopF1e, and HopF4a from *P. syringae* pv. *actinidiae* [Psa] V-13; and HopF1b from *P. syringae* pv. *actinidifoliorum* [Pfm] LV-5, aligned with ClustalW.

**Table S1.**
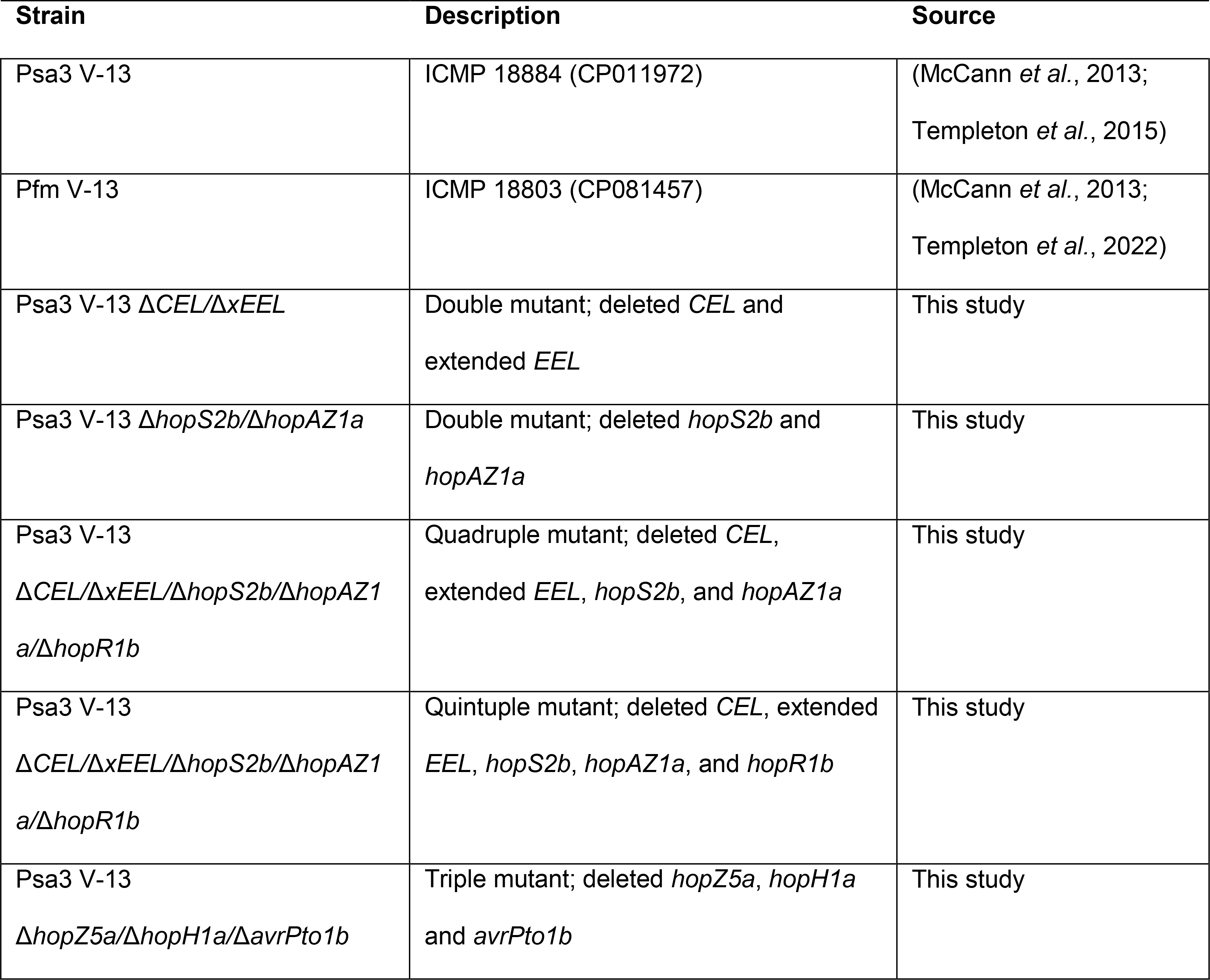
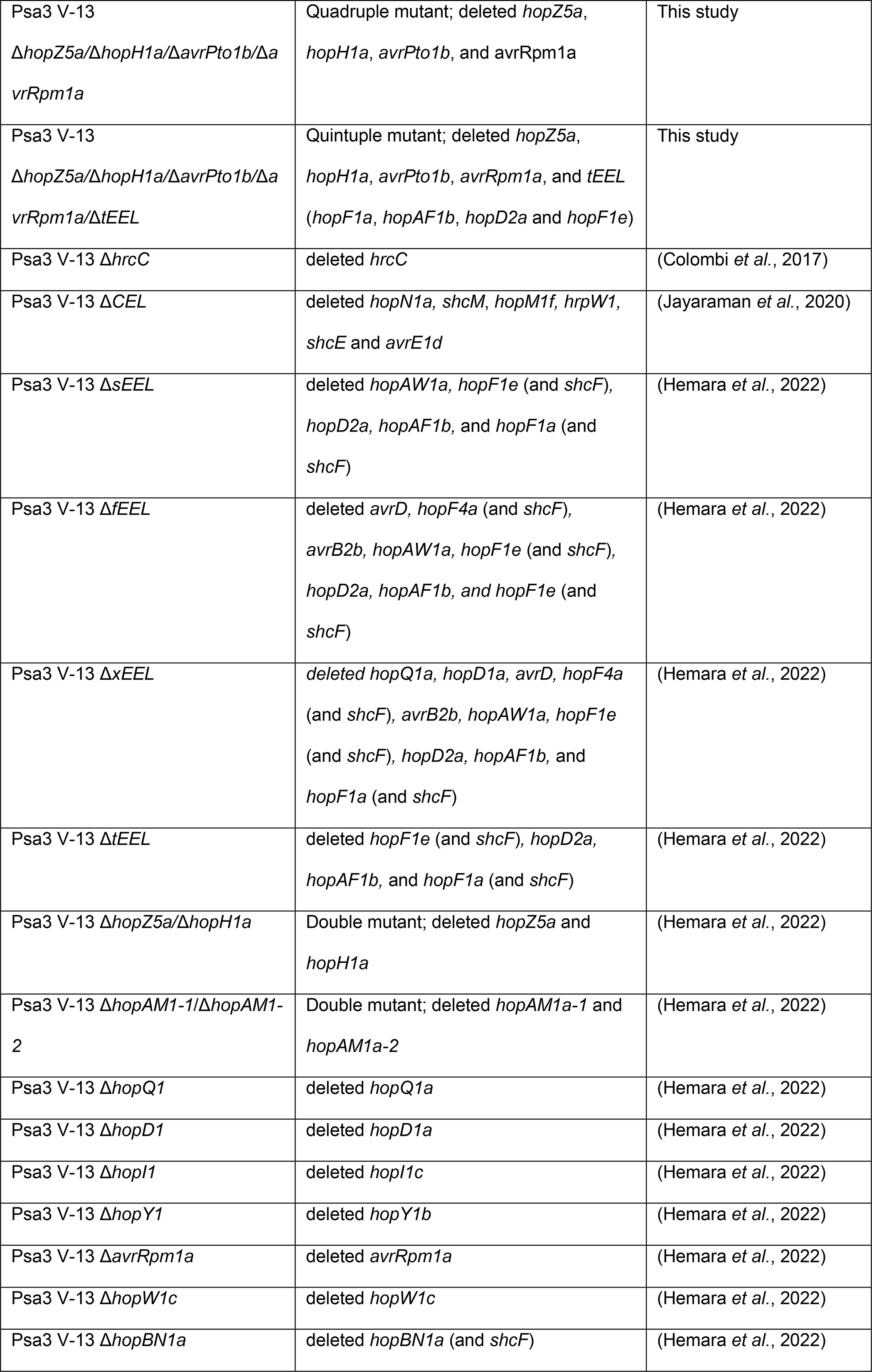
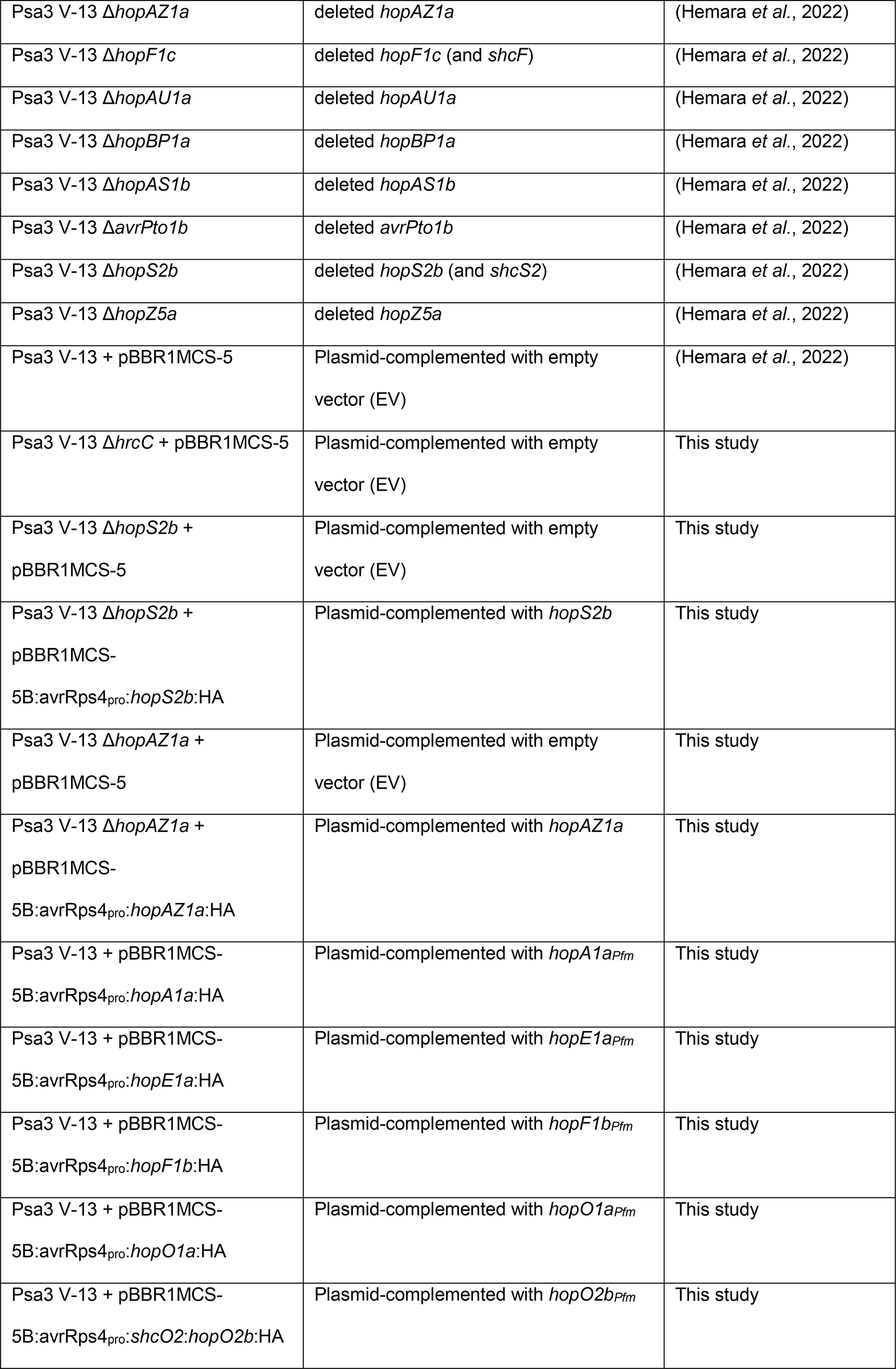
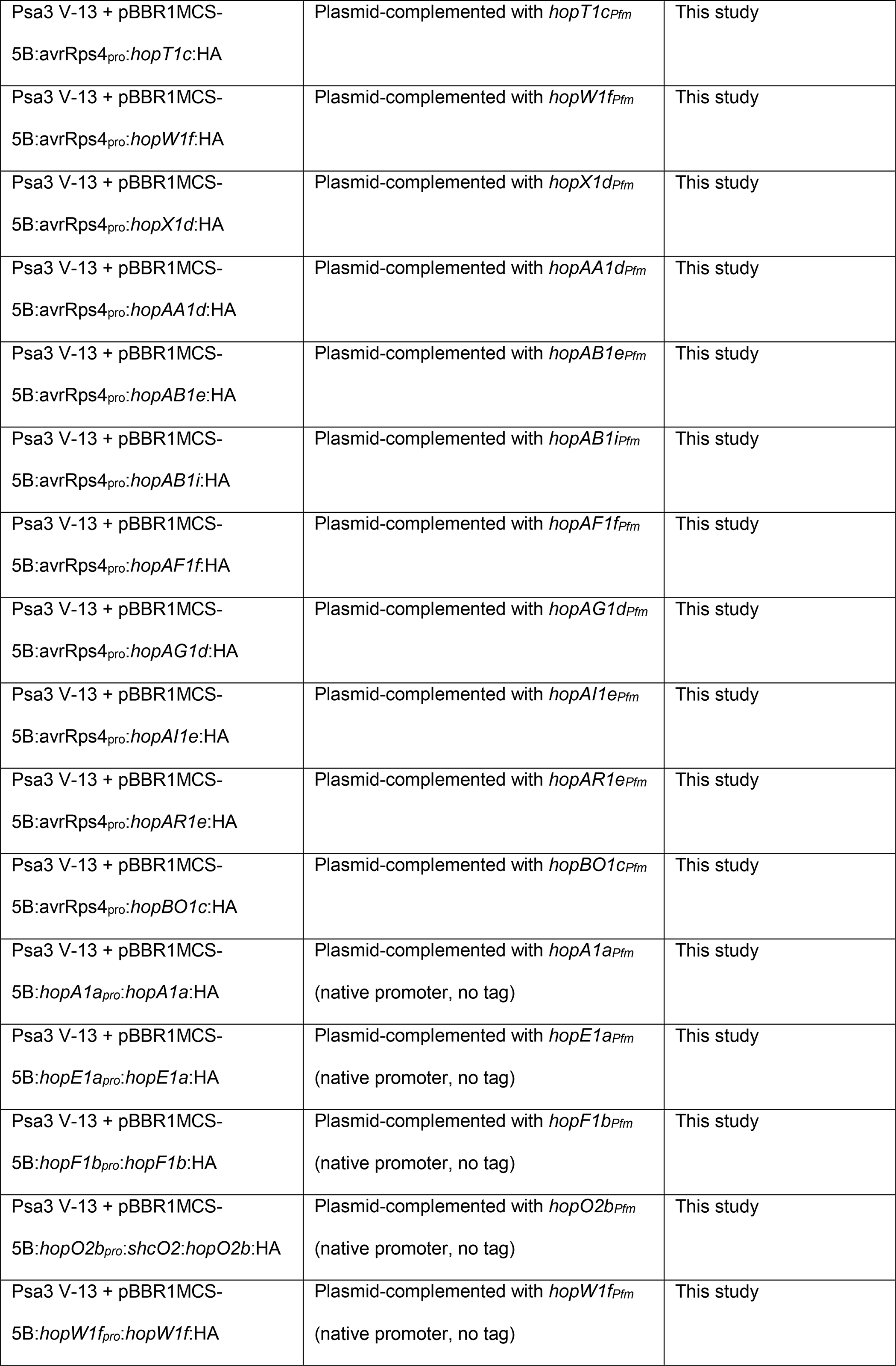
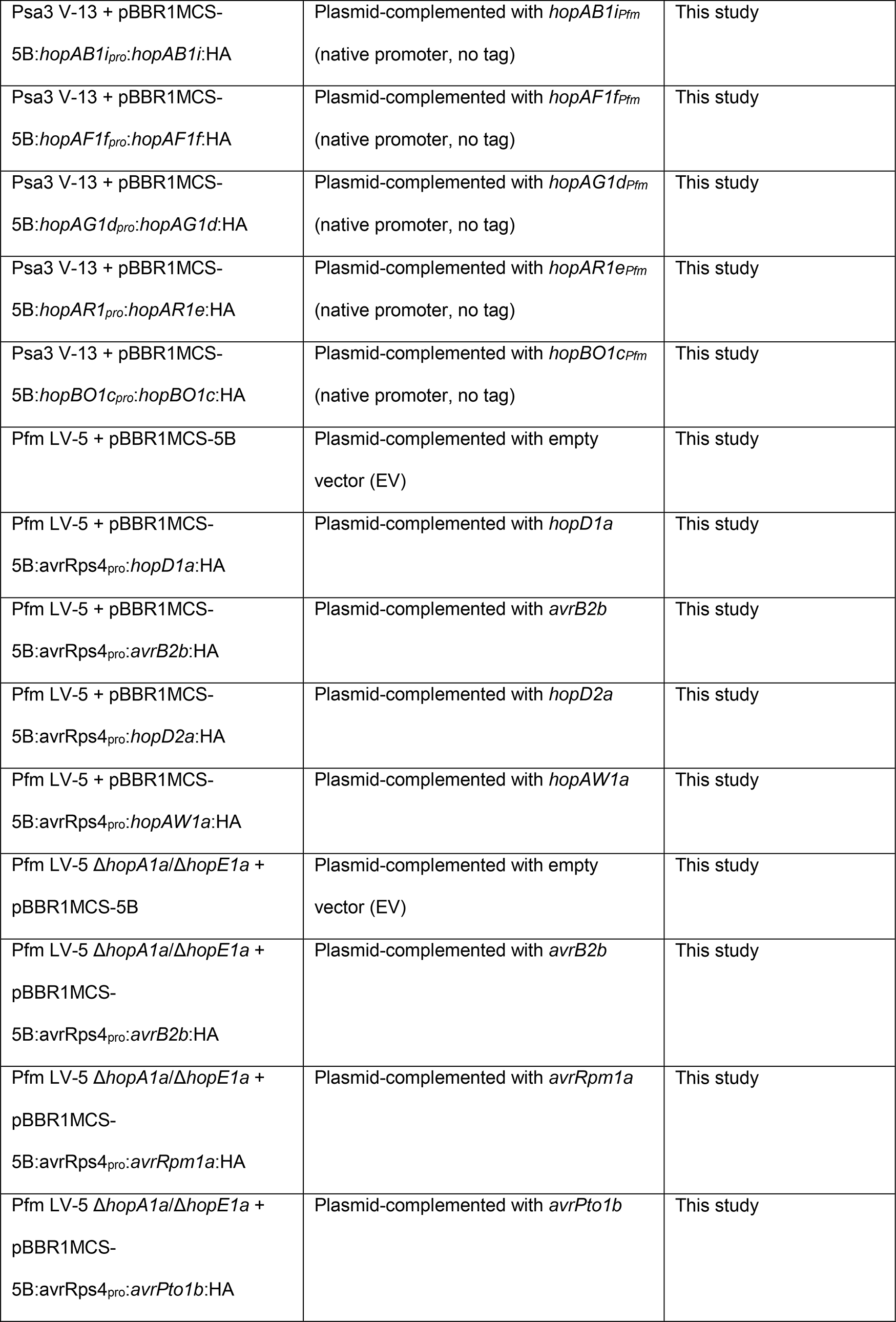
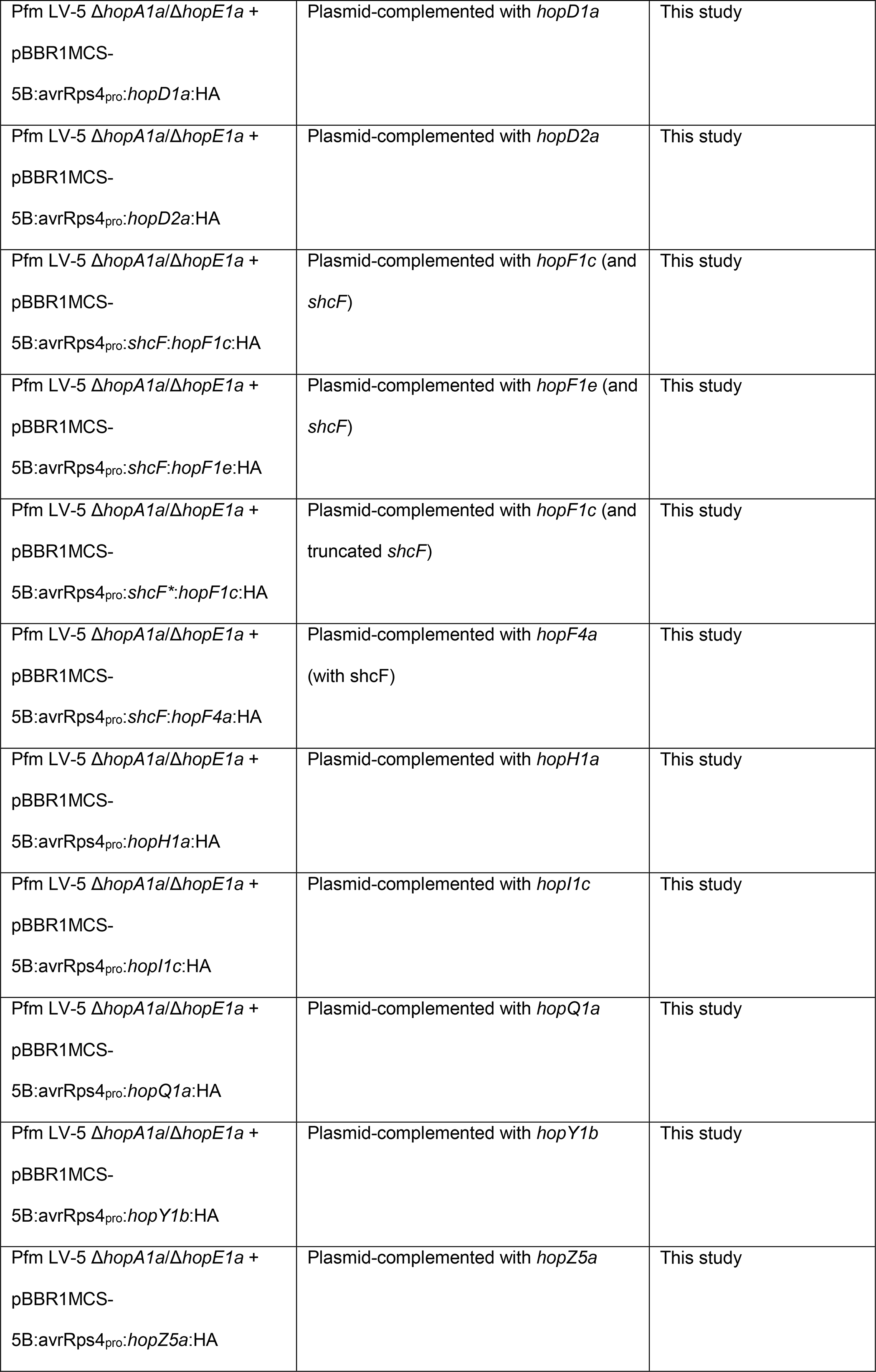
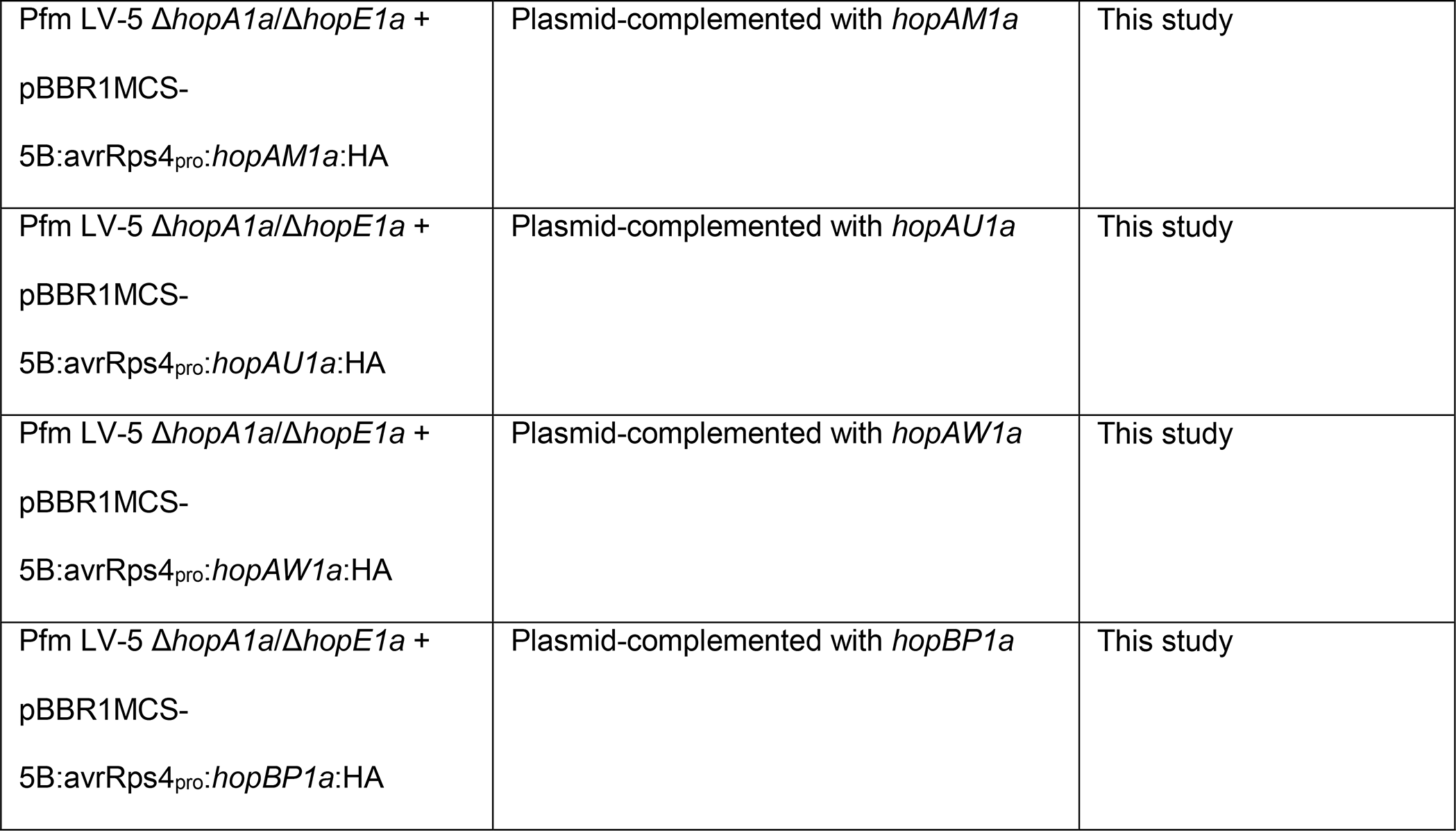
Pseudomonas syringae pv. actinidiae and P. syringae pv. actinidifoliorum strains used in this study.

**Table S2.**
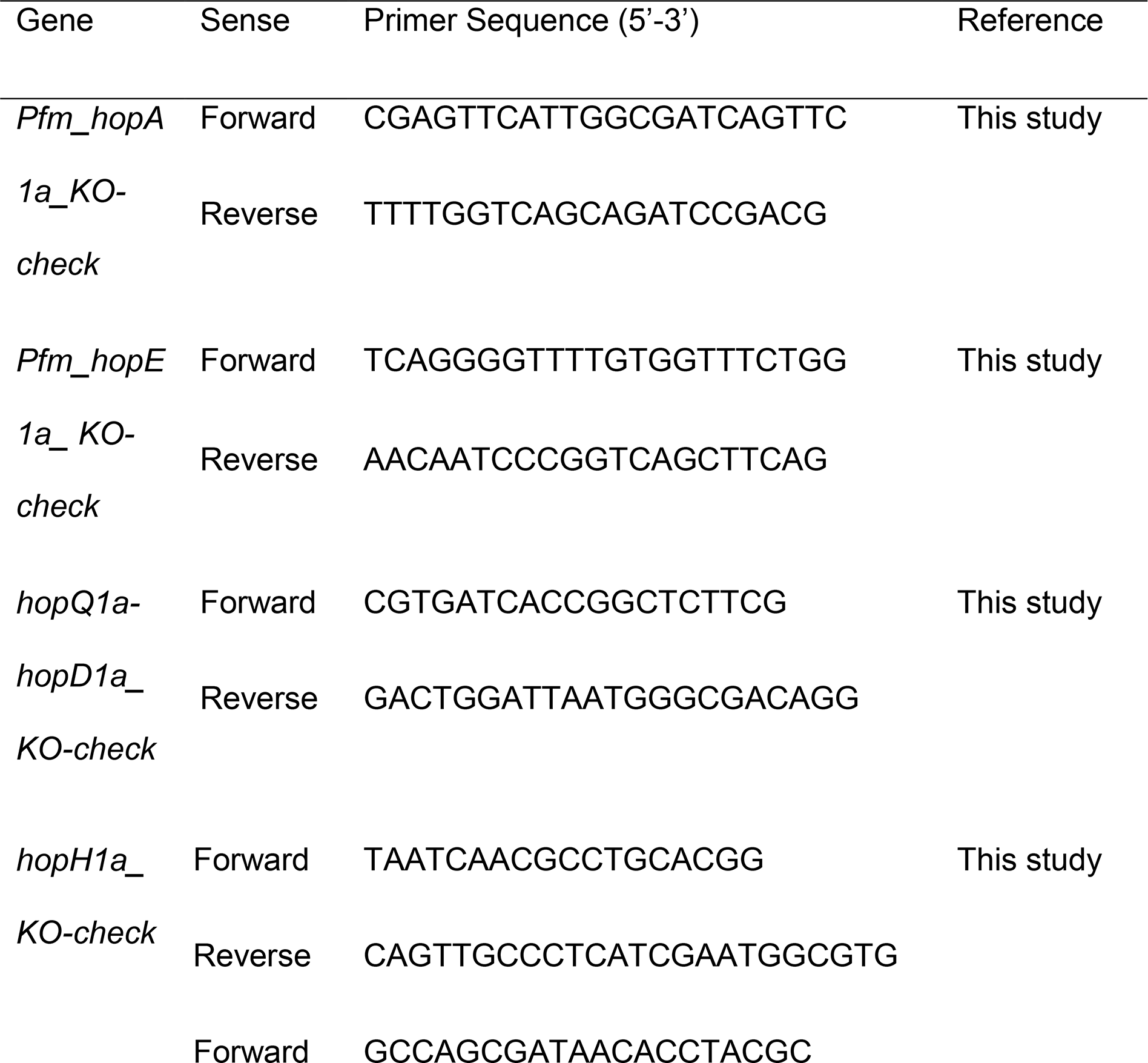
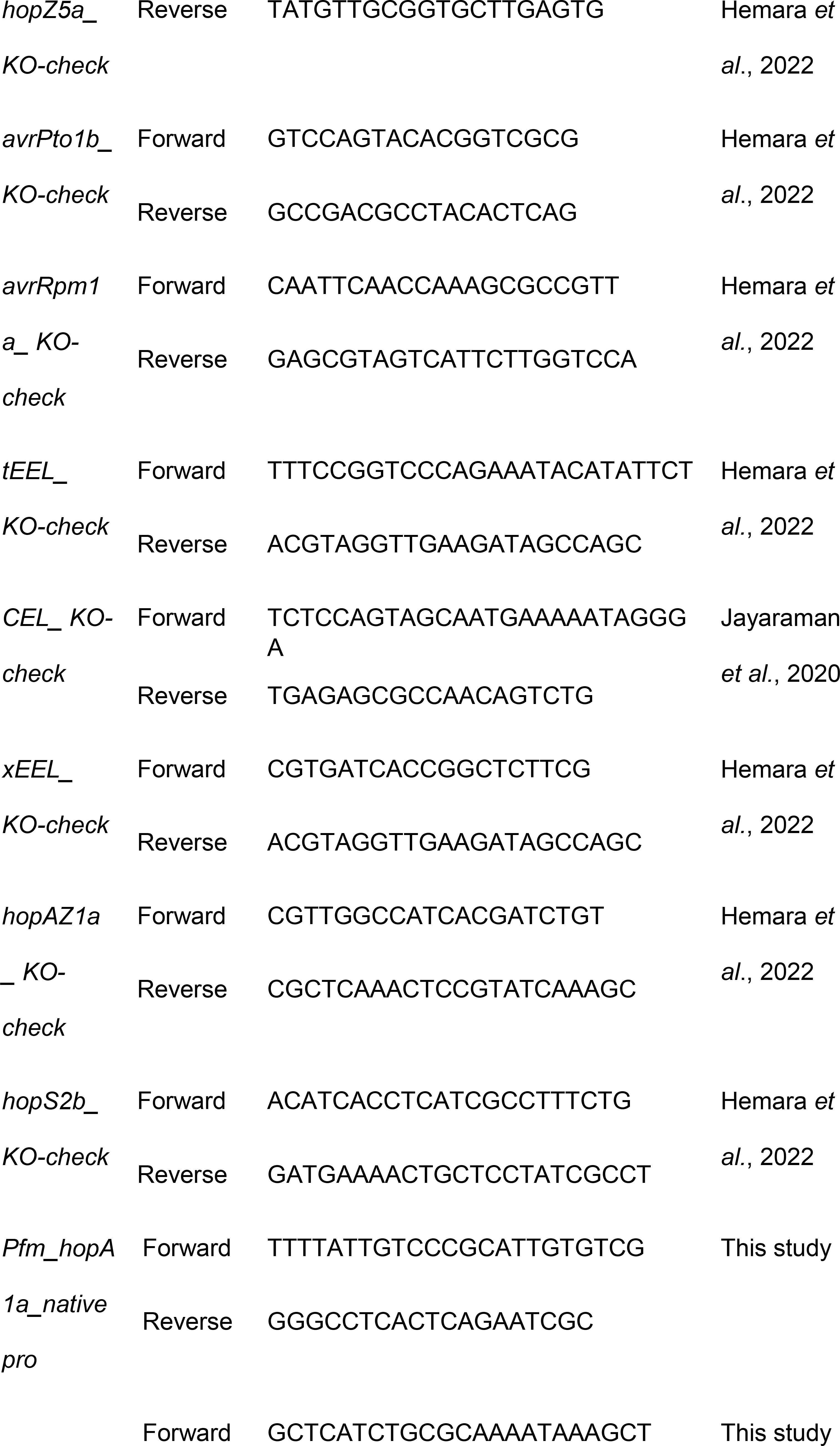
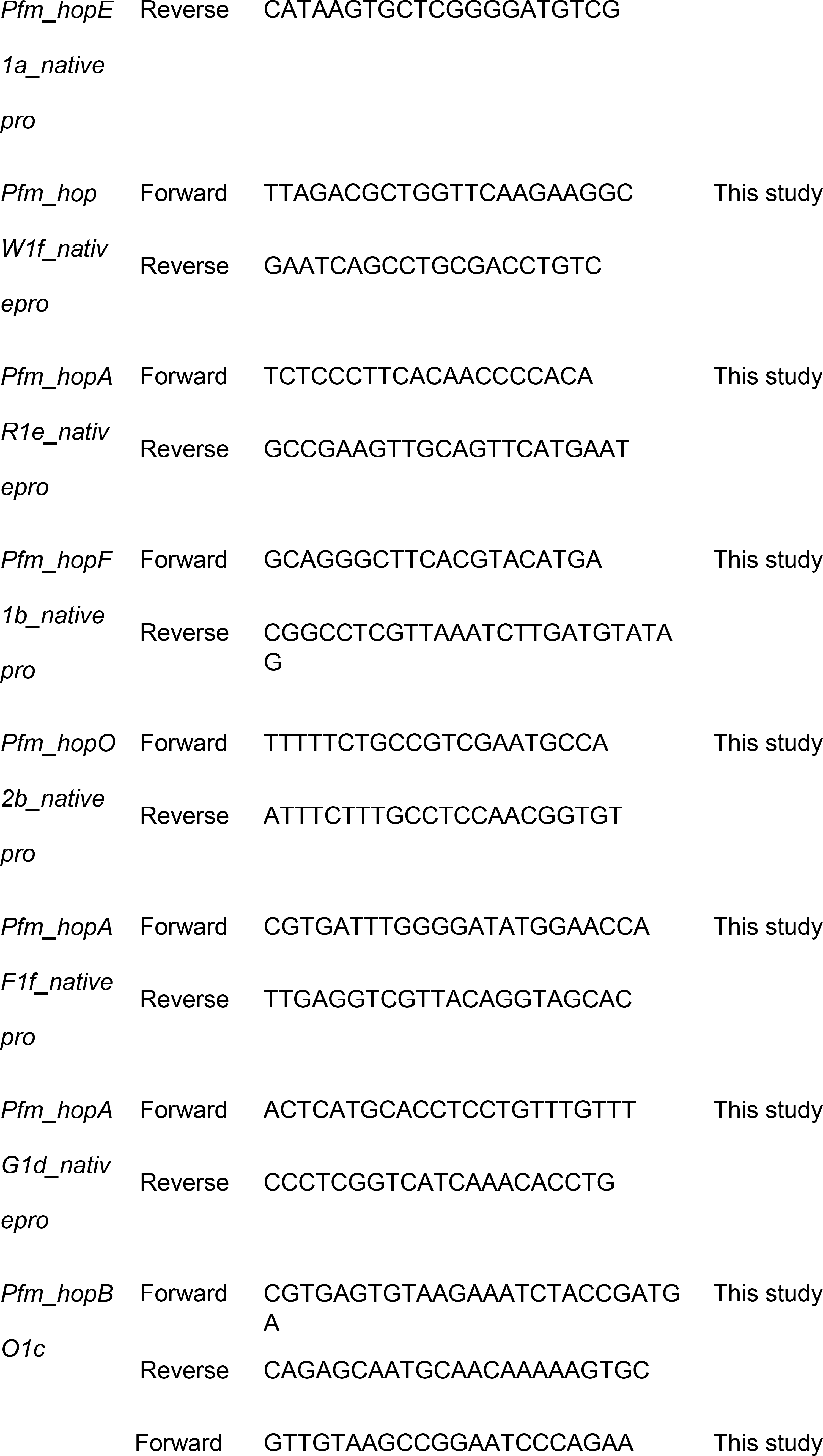
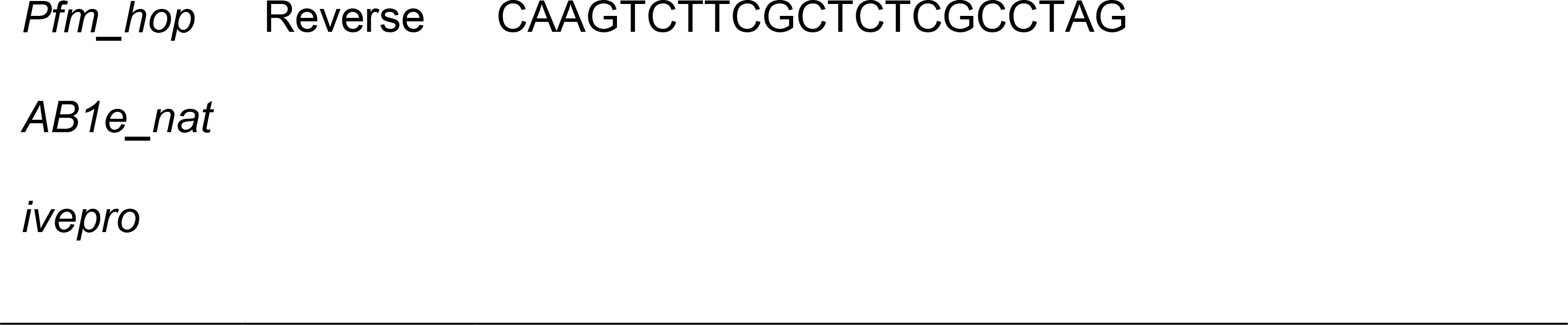
Primers used in this study.

Suppl Table S3
| Effectors | *Psa3* V-13 | *Pfm* LV-5 | AA identity (%) |
| --- | --- | --- | --- |
| AvrB2b | o |  |  |
| AvrE1d | o | o | 97 |
| AvrPto1b | o |  |  |
| AvrRpm1a | o |  |  |
| HopA1a |  | o |  |
| HopD1a | o |  |  |
| HopE1a |  | o |  |
| HopF1a | o |  |  |
| HopF1b |  | o |  |
| HopF1c | o |  |  |
| HopH1a | o |  |  |
| HopZ5a | o |  |  |
| HopI1c | o |  |  |
| HopM1f | o | o | 99 |
| HopN1a | o | o | 99 |
| HopO1a |  | o |  |
| HopQ1a | o |  |  |
| HopR1b | o | o | 99 |
| HopO2b |  | o |  |
| HopS2b/HopS2c | o | o | 95 |
| HopT1c |  | o |  |
| HopW1f |  | o |  |
| HopX1d |  | o |  |
| HopBO1c |  | o |  |
| HopF4a | o |  |  |
| HopY1b | o |  |  |
| HopBP1a | o |  |  |
| HopAA1d |  | o |  |
| HopAB1i |  | o |  |
| HopW1c | o | o | 99 |
| HopAF1b/f | o | o |  |
| HopAF1f |  | o |  |
| HopAG1d |  | o |  |
| HopAH1a | o | o | 99 |
| HopAH1j/i | o | o | 94 |
| HopAH1k/c | o | o | 88 |
| HopAI1b/e | o | o | 82 |
| HopAM1a-1/-2 | o |  |  |
| HopD2a | o |  |  |
| HopAR1e |  | o |  |
| HopAS1b | o | o | 99 |
| HopAU1a | o |  |  |
| HopAW1a | o |  |  |
| HopAB1e |  | o |  |
| HopAZ1a | o | o | 96 |
| HopF1e | o |  |  |
| HopBN1a | o |  |  |

Red = *Psa3* V-13 effectors required for virulence Green = *Pfm* LV-5 unique effectors Blue = *Psa3* V-13 unique effectors

